# Dietary restriction and *clock* delay eye aging to extend lifespan in *D. melanogaster*

**DOI:** 10.1101/2021.05.08.443272

**Authors:** Brian A. Hodge, Geoffrey T. Meyerhof, Subhash D. Katewa, Ting Lian, Charles Lau, Sudipta Bar, Nicole Leung, Menglin Li, David Li-Kroeger, Simon Melov, Birgit Schilling, Craig Montell, Pankaj Kapahi

**Affiliations:** Buck Institute for Research on Aging, 8001 Redwood Blvd, Novato, CA 94945, USA; NGM Biopharmaceuticals, 333 Oyster Point Blvd, South San Francisco, CA 94080, USA; Sichuan Agricultural University, 46 Xinkang Rd, Yucheng District, Ya’an, Sichuan, China; Neuroscience Research Institute and Department of Molecular, Cellular and Developmental Biology, University of California, Santa Barbara, Santa Barbara, 93106, California, USA; Department of Psychiatry and Behavioral Sciences, Stanford University, Stanford, CA 94305, USA; Department of Neurobiology, Stanford University, Stanford, CA 94305, USA; Department of Neurology, Baylor College of Medicine, Houston, TX 77096, USA

## Abstract

Many vital processes in the eye are under circadian regulation, and circadian dysfunction has emerged as a potential driver of eye aging. Dietary restriction is one of the most robust lifespan-extending therapies and amplifies circadian rhythms with age. Herein, we demonstrate that dietary restriction extends lifespan in *D. melanogaster* by promoting circadian homoeostatic processes that protect the visual system from age- and light-associated damage. Disrupting circadian rhythms in the eye by inhibiting the transcription factor, Clock (CLK), or CLK-output genes, accelerated visual senescence, induced a systemic immune response, and shortened lifespan. Flies subjected to dietary restriction were protected from the lifespan-shortening effects of photoreceptor activation. Inversely, photoreceptor inactivation, achieved via mutating rhodopsin or housing flies in constant darkness, primarily extended lifespan in flies reared on a high-nutrient diet. Our findings establish the eye as a diet-sensitive modulator of lifespan and indicate that vision is an antagonistically pleiotropic process that contributes to organismal aging.

## Introduction

Circadian rhythms are approximate 24-hour oscillations in behavior, cellular physiology, and biochemistry, which evolved to anticipate and manage predictable changes associated with the solar day (e.g., predator/prey interactions, nutrient availability, phototoxicity, etc.) [1]. Circadian rhythms are generated by endogenous clocks that sense time-cues (e.g., light and food) to govern rhythmic oscillations of gene transcriptional programs, synchronizing cellular physiology with daily environmental stressors [2]. In addition to keeping time, the molecular clock regulates the temporal expression of downstream genes, known as clock-controlled genes, to promote tissue-specific rhythms in physiology [3]. The *Drosophila* molecular clock is comprised of transcriptional-translational feedback loops, where the transcription factors Clock (CLK) and Cycle (CYC) rhythmically activate their own repressors, Period and Timeless [2]. This feedback loop not only exists in central pacemaker neurons, where it sets rhythms in locomotor activity, it also functions in peripheral tissues, such as the eye [4].

Aging is associated with a progressive decline in visual function and an increase in the incidence of ocular disease. *Drosophila* photoreceptor cells serve as a powerful model of both visual senescence and retinal degeneration [5, 6]. *Drosophila* and mammalian photoreceptors possess a cell-intrinsic molecular clock mechanism that temporally regulates a large number of physiological processes, including light-sensitivity, metabolism, pigment production, and susceptibility to light-mediated damage [7]. Visual senescence is accompanied by a reduced circadian amplitude in core-clock gene expression within the retina [8]. This reduction in retinal circadian rhythms may be causal in eye aging, as mice harboring mutations in their core-clock genes, either throughout their entire body, or just in their photoreceptor cells, display several early-onset aging phenotypes within the eye. These mice prematurely form cataracts and have reduced photoreceptor cell light-sensitivity and viability [8]. However, the molecular mechanisms by which the molecular clock influences eye aging are not fully understood.

Dietary restriction (DR), defined by reducing specific nutrients or total calories, is the most robust mechanism for delaying disease and extending lifespan [9]. The mechanisms by which DR promotes health and lifespan may be integrally linked with circadian function, as DR enhances the circadian transcriptional output of the molecular clock and preserves circadian function with age [10]. Inversely, high-nutrient diets (i.e., excess consumption of protein, fats, or total calories) repress circadian rhythms and accelerate organismal aging [11, 12]. However, how DR modulates circadian rhythms within the eye, and how these rhythms influence DR-mediated lifespan extension, had yet to be examined.

Herein, we sought to elucidate the circadian processes that are activated by DR by performing an unbiased, 24-hour time-course mRNA expression analysis in whole flies. We found that circadian processes within the eye are highly elevated in expression in flies reared on DR. In particular, DR enhanced the rhythmic expression of genes involved in the adaptation to light (i.e., calcium handling and de-activation of rhodopsin-mediated signaling). Building on this observation, we demonstrate that the majority of these circadian phototransduction components were transcriptionally regulated by CLK. Eliminating CLK function either pan-neuronally, or just in the photoreceptors, accelerated visual decline with age. Furthermore, disrupting photoreceptor homeostasis increased systemic immune responses and shortened lifespan. Several eye-specific CLK-output genes that were upregulated in expression in response to DR, were also required for DR- to slow visual senescence and extend lifespan.

## Results

### Dietary restriction amplifies circadian transcriptional output and delays visual senescence in a CLK-dependent manner

To determine how DR changes circadian transcriptional output, we performed a series of microarray experiments over the span of 24-hours in female *Canton-S* flies (whole body) reared on either a high-yeast (5%; *ad libitum*, AL) diet or a low-yeast (0.5%; DR) diet (**Supplementary Fig. 1a**). Flies maintained on DR displayed nearly twice the number circadian transcripts compared to flies on AL (**Fig. 1a, b and Supplementary Fig. 1b**). Circadian gene expression was also more robust on DR vs AL. DR-specific oscillators were statistically more rhythmic (lower JTK_CYCLE circadian *p*-values) and displayed larger circadian amplitudes than AL-specific oscillators (**Supplementary Fig. 1c, d**). Diet also drastically altered the circadian transcriptional profile, as only 16% of DR oscillators were also oscillating on AL (**Supplementary Fig. 1b**). Furthermore, the AL and DR circadian transcriptomes were enriched for distinct processes (**Supplementary Fig. 1e, f and Supplementary Data 1**).

**Figure 1.**
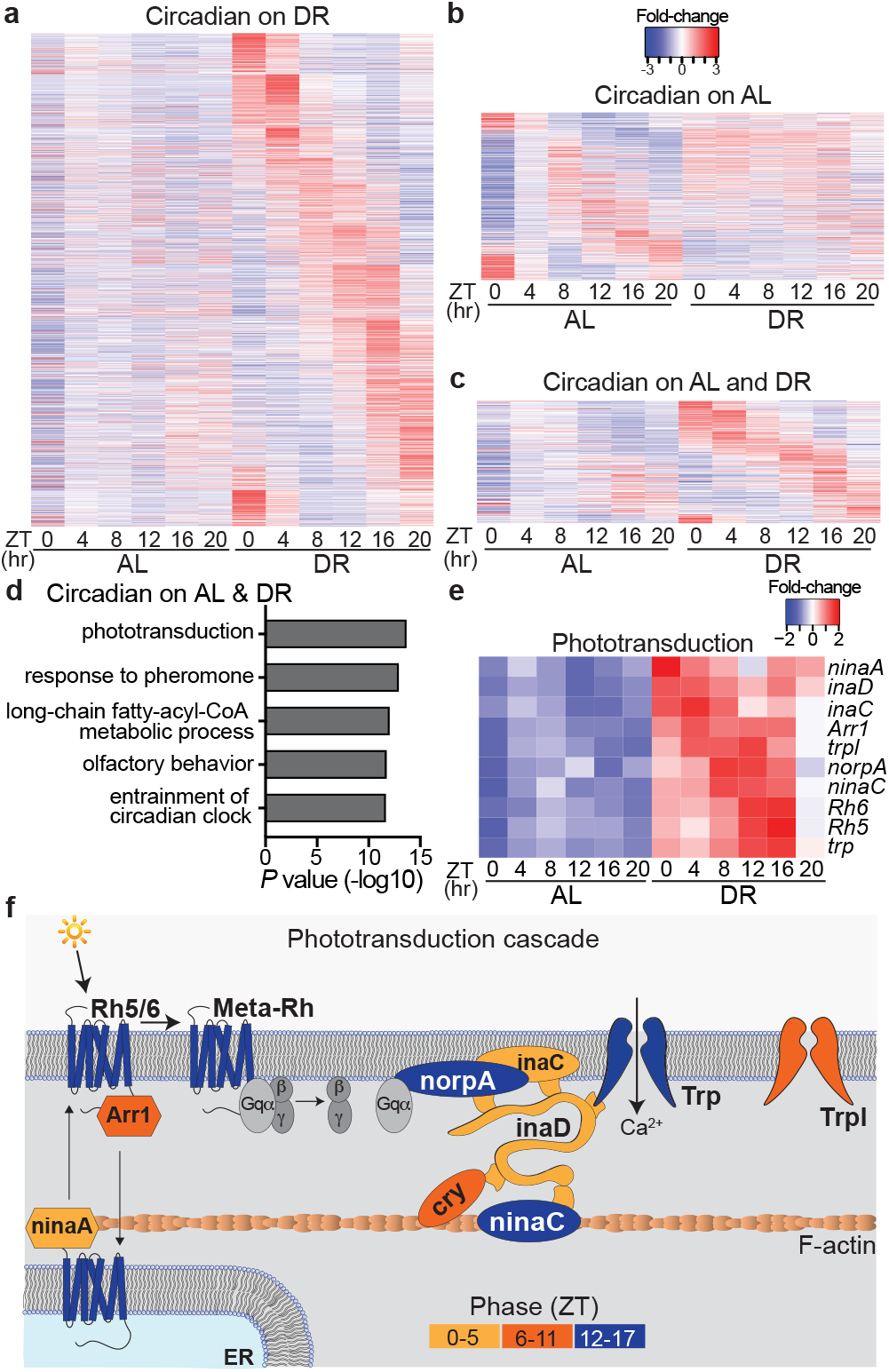
Dietary restriction amplifies circadian transcriptional output and rhythmicity of phototransduction genes. (**a-c**) Circadian transcriptome heatmaps for Canton-S flies representing 24-hour expression plots for transcripts that cycle only on DR (a, *n*=1609 transcripts), only on AL (b, *n*=568 transcripts), or on both diets (c, *n*=301 transcripts). Circadian transcripts (24h period, JTK_CY- CLE *p*value<0.05) are plotted by phase. (**d**) Gene-ontology enrichment categories corresponding to transcripts that cycle on both AL and DR diets. (**e**) Heatmap of phototransduction transcript expression on AL and DR. (**f**) Phototransduction cascade diagram with components colored according to their circadian phase on DR.

Transcripts that oscillate on both AL and DR diets were highly enriched for genes that comprise the canonical phototransduction signaling cascade (**Fig. 1c-d**), which is the process by which *Drosophila* photoreceptor cells, the primary light-sensitive neurons, transduce light information into a chemical signal [13]. Briefly, light-mediated conversion of rhodopsin proteins to their meta-rhodopsin state stimulates heterotrimeric Gq proteins that activate phospholipase C (*norpA*), which produces secondary messengers and promotes the opening of Transient Receptor Potential channels (TRP, TRPL), ultimately allowing Ca^2+^ and Na^+^ to depolarize the photoreceptor cell [14]. Although the phototransduction transcripts were cyclic on both diets, on DR their expression became more rhythmic (lower JTK_CYCLE *p*-values & larger circadian amplitudes) and elevated (∼2-fold increase in expression across all timepoints) (**Fig. 1e and Supplementary Fig. 1i**). Since our time-course analyses were perfromed in whole-fly, we queried publicly available circadian transcriptomes from wild-type heads to further investigate the rhythmic oscillations of eye-related transcripts [15]. The majority of the DR-sensitive phototransduction genes also robustly cycled in wild-type heads (**Supplementary Table 1**). Furthermore, the GO-term “phototransduction” (GO:0007602) was amongst the most enriched cyclic processes in the heads of wild-type flies, as ∼70% of the genes that comprise the category oscillate in a circadian fashion (**Supplemental Data 2**).

In *Drosophila* and mammals, visual function oscillates to align with daily changes in ambient illuminance from the sun, which can be 10^6^ to 10^8^-fold brighter during the day than at night [16]. Photoreceptors are unique in that they have evolved mechanisms responsible for maintaining homeostasis in the presence of light-induced calcium ion gradients that are magnitudes greater than what other neuronal populations experience [17, 18]. Mechanisms of light adaptation within photoreceptors include the rapid (millisecond) closure of TRP channels (facilitated via enzymes scaffolded by *inaD*), rhodopsin internalization from the rhabdomere membrane (e.g., *arr1*, *arr2*), and calcium efflux (e.g., *calx*) [19, 20]. Acrophase analyses (i.e., time of peak expression) revealed that circadian transcripts that promote photoreceptor activation (Ca^2+^ influx) reach peak expression during the dark-phase, while genes that terminate the phototransduction response (i.e., deactivation of rhodopsin mediated signaling) peak in anticipation of the light-phase (**Fig. 1f and Supplementary Fig. 1j**). These findings provide a potential mechanistic explanation for the rhythmic response pattern in light-sensitivity observed in *Drosophila* photoreceptors and suggests that DR’s ability to delay visual senescence is mediated in part by amplifying circadian rhythms within photoceptors (See **Supplemental Discussion 1** for additional interpretations).

To determine if molecular clocks mediate the enhanced rhythmic expression of phototransduction genes on DR, we measured the transcriptome of fly heads with pan-neuronal over-expression of a dominant negative form of the core-clock factor, CLK (Elav-GS-GAL4>UAS-CLK-Δ1; denoted nCLK-Δ1) (**Supplementary Fig. 2a**). To avoid potential developmental defects related to Clk disruption, we used a drug-inducible (RU486) “gene-switch” driver to express CLK-Δ in adult flies. Genes downregulated in nCLK-Δ1 heads were enriched for light-response pathways, including “response to light stimulus” and “deactivation of rhodopsin signaling” (**Fig. 2a, b and Supplemental Data 3**). Additionally, genes that were both circadian in wild-type heads and downregulated in nCLK-Δ1 were highly enriched for homeostatic processes related to eye function (**Supplementary Fig. 2c and Supplemental Data 4**). Together, this indicates that CLK governs the circadian transcriptional regulation of many eye-related processes in *Drosophila*.

**Figure 2.**
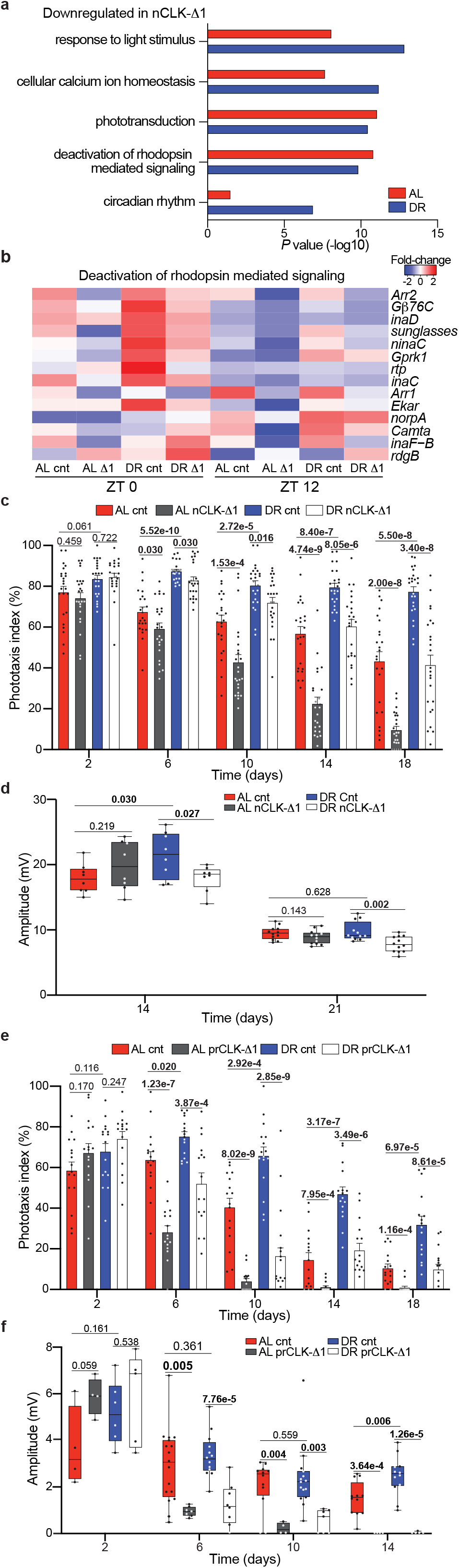
Dietary restriction delays visual senescence in a CLK-dependent manner. (**a**) GO enrichment scores corresponding to downregulated light-response genes in heads from RNA-Seq of nCLK-Δ1 (Elav-GS-GAL4> UAS-CLK-Δ1) vs controls. (**b**) Heatmap of normalized RNA-Seq expression corresponding to the gene-ontology category “Deactivation of rhodopsin mediated signaling” (GO:0016059) in nCLK-Δ1 and controls at zeitgeber times 0 and 12 (lights on and lights off, respectively). (**c**) Positive phototaxis responses for nCLK-Δ1 flies. For each timepoint results are represented as average percent positive phototaxis +/- SEM (*n*=24 biological replicates, *N*=480 flies per condition). (**d**) Boxplots of electroretinogram amplitudes for nCLK-Δ1 flies and controls at day 14 and 21. (e) Positive phototaxis responses for prCLK-Δ1 flies (Trpl-GAL4; GAL80^ts^> UAS-CLK-Δ1) and control flies (Trpl-GAL4; GAL80^ts^>*CantonS*) reared at 30°C. For each timepoint results are represented as average percent positive phototaxis +/- SEM (*n*=16 biological replicates, *N*=320 flies per condition). (**f**) Boxplots of electroretinogram amplitudes for prCLK-Δ1 and control flies reared at 30°C. Illuminance was set at 150 Lux. (c-f) *P*values were determined by two-tailed Student’s *t*test (unpaired), comparing responses between diet and/or genotype at each timepoint.

Given DR’s ability to improve homeostasis across an array of tissues [21], and its ability to enhance the circadian rhythmicity of light-response genes, we examined how diet and clocks influence visual function with age. We longitudinally quantified the positive phototaxis response of wild-type flies (*Canton-S* and *Oregon-R*) reared on either AL or DR diets (experimental setup in Supplementary Fig. 2d). Compared to AL-fed flies, DR slowed the decline in positive phototaxis observed with age (**Supplementary Fig. 2e, f**). Importantly, this effect cannot solely be attributed to diet-dependent changes in locomotor activity, as climbing activity and phototaxis declined at different rates with age (**Supplementary Fig. 2g**). Compared to wild-type flies, DR minimally protected *Clk^out^* (*Clk*-null) flies from age-related declines in phototaxis (**Supplementary Fig. 2h**). nCLK-Δ1 and nCLK-Δ2 (an additional dominant negative *Clk* mutant, Elav-GS-GAL4>UAS-CLK-Δ2) flies displayed accelerated declines in positive phototaxis with age compared to controls (**Fig. 2c and Supplementary Fig. 2i**). Since the positive phototaxis assay measures a behavioral response to light, we next evaluated how diet and CLK directly influence photoreceptor function with age by performing extracellular electrophysiological recordings of the eye (electroretinograms, ERG [22]). We observed larger ERG amplitudes, i.e. the light-induced summation of receptor potentials from the photoreceptors [23], in control flies reared on DR vs AL at day 14 (**Fig. 2d**). Furthermore, the DR-mediated enhancements in the ERG amplitudes were significantly reduced in nCLK-Δ1 flies with age (**Fig. 2d**).

Since the Elav-GS-GAL4 driver is expressed in a pan-neuronal fashion (i.e., photoreceptors + extra-ocular neurons), we sought to examine how disrupting CLK function solely within photoreceptors influences visual function with age. To this end, we crossed UAS-CLK-Δ1 flies with a photoreceptor-specific GAL4 driver line under the temporal control of the temperature sensitive GAL80 protein (Trpl-GAL4; GAL80^ts^>UAS-CLK-Δ1, denoted prCLK-Δ1). To avoid disrupting CLK function during development, prCLK-Δ1 flies were raised at 18°C (GAL80 active, GAL4 repressed) and then transferred to 30°C (GAL80 repressed, GAL4 active) following eclosion. When compared to control flies (Trpl-GAL4; GAL80^ts^>*CantonS*), prCLK-Δ1 flies displayed accelerated declines in both positive phototaxis and ERG amplitude with age, and in a similar fashion to the nCLK-Δ1 flies (**Fig. 2e-f**). Together, our gene expression, phototaxis, and ERG data indicate that DR functions in a CLK-dependent manner to delay photoreceptor aging in the fly.

### nCLK-Δ drives a systemic immune response and reduces longevity

Age-related declines in tissue homeostasis are accompanied by elevated immune responses and inflammation [24, 25]. Interestingly, we found that genes upregulated in nCLK-Δ1 fly heads were significantly enriched for immune and antimicrobial humoral responses (**Fig. 3a, b**). In *Drosophila,* damage-associated molecular patterns can induce a sterile immune response that is characterized by the expression of anti-microbial peptides (AMPs), similar to the effects from infections by pathogens [26]. We quantified the mRNA expression of AMPs in the bodies of nCLK-Δ1 and nCLK-Δ2 flies to determine if neuronal damage signals propagate throughout the body to drive systemic immune responses; the *Drosophila* fat body generates high levels of AMPs in response to intrinsic damage signals [26]. AMP expression (*AttA*, *DiptB*, and *Dro*) was reduced in control flies reared on DR compared to AL, however nCLK-Δ1 and nCLK-Δ2 elevated AMP expression on DR (**Fig. 3c and Supplementary Fig. 3a**). To further investigate this systemic inflammatory response, we isolated and quantified hemolymph from nCLK-Δ1 and control flies. In agreement with the transcriptional activation of AMPs in both the heads and bodies of nCLK-Δ1 flies, we found the most highly upregulated protein in nCLK-Δ1 hemolymph to be the antimicrobial peptide, AttC (Supplementary Fig. 3b). Furthermore, we observed an enrichment for proteins associated with translational activation (e.g., cytoplasmic translation and ribosomal biogenesis) within the upregulated proteins in the nCLK-Δ1 hemolymph, which may reflect the activation of hemocytes, the immune effector cells in *Drosophila* (**Supplemental Data 5**) [27]. Taken together, these data demonstrate that disrupting neuronal CLK function elevates systemic immune responses.

**Figure 3.**
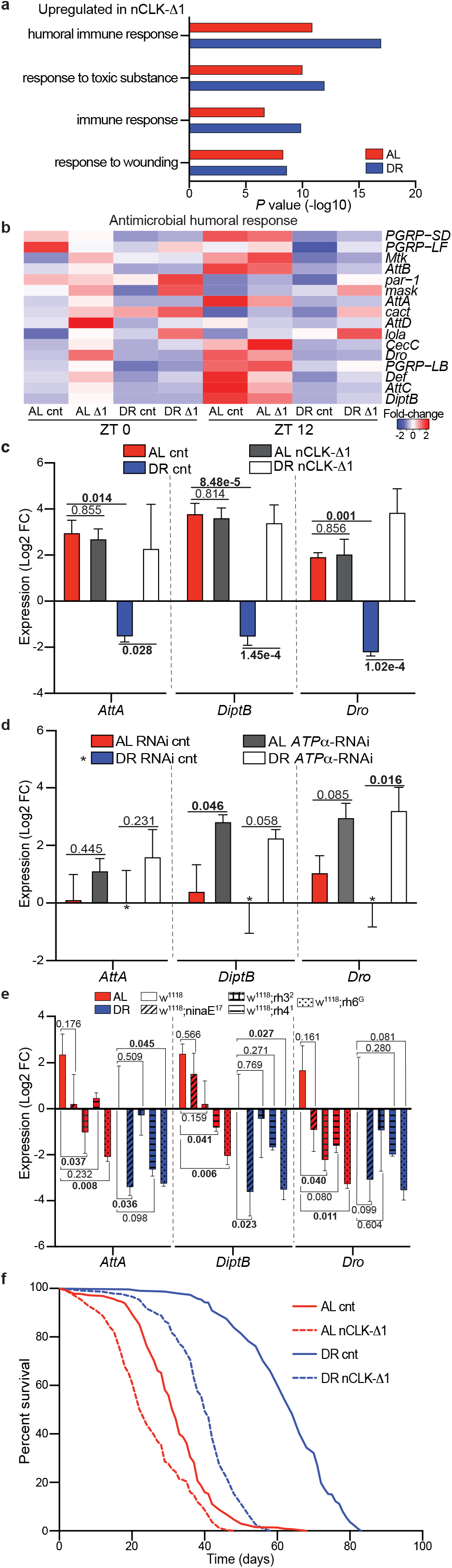
nCLK-Δ1 flies display elevated immune responses and shortened lifespan. (**a**) GO enrichment scores corresponding to upregulated inflammatory genes in heads from RNA-Seq of nCLK-Δ1 vs controls. (**b**) Heatmap of normalized RNA-Seq expression corresponding to the gene-ontology category “Antimicrobial humoral responses” (GO:0019730) in nCLK-Δ1 and controls. (**c**) Relative expression of AMP genes (*AttA*, *DiptB*, and *Dro*) calculated by RT-qPCR with mRNA isolated from nCLK-Δ1 bodies. Results are plotted as average Log2 fold-change in expression calculated by the ΔΔ-Ct method, normalized to DR vehicle treated control samples, as well as the housekeeping gene *rp49* +/- SEM (*n*=3 biological replicates, *N*=30 flies per biological replicate). (**d**) Relative mRNA expression of AMP genes calculated by RT-qPCR with mRNA isolated from bodies of eye-specific *ATP*α knockdown flies (GMR-GAL4>UAS-*ATP*α-RNAi) vs RNAi control flies (GMR-GAL4>UAS-*mCherry*-RNAi). Results are plotted as average Log2 fold-change in expression calculated by the ΔΔ-Ct method, normalized to DR RNAi control samples as well as housekeeping gene *rp49* +/- SEM (*n*=3 biological replicates, *N*=30 flies per biological replicate). (**e**) Relative mRNA expression of immune genes (*AttaA*, *DiptB*, and *Dro*) calculated by RT-qPCR with mRNA isolated from bodies of w1118 and rhodopsin mutant flies housed in 12:12h LD. Results are plotted as average Log2 fold-change in expression calculated by the ΔΔ-Ct method normalized *w^1118^* DR control samples as well as *rp49* +/- SEM (*n*=3 biological replicates, *N*=30 flies per biological replicate). (**f**) Kaplan-Meyer survival analysis of nCLK-Δ1 flies (Elav-GS-GAL4>UAS-CLK-Δ1). Survival data is plotted as an average of three independent lifespan repeats. Control flies (vehicle treated): AL *N*=575, DR *N*=526; nCLK-Δ1 flies (RU486 treated): AL *N*=570, DR *N*=565. (c-e) *P*values were calculated with the pairwise Student’s *t*test comparing Log2 fold-changes in expression.

To determine if photoreceptor degeneration induces a systemic immune response in *Drosophila*, we forced photoreceptor degeneration by knocking down *ATPα* within the eye (GMR-GAL4>UAS-*ATPα*-RNAi), and quantified expression of AMPs within the bodies. *ATPα* encodes the catalytic alpha subunit of the Na^+^K^+^ATPase responsible for reestablishing ion balance in the eye during light responses [28, 29]. Our decision to use *ATPα* knockdown as a model of photoreceptor degeneration was motivated by previous reports indicating that its expression is under circadian regulation [30] and that its knockdown in the eye results in aberrant ion homeostasis that drives age-dependent, light-independent photoreceptor degeneration [31]. Ocular knockdown of *ATPα* rendered flies blind in both AL and DR conditions compared to controls (**Supplementary Fig. 3c**). Knocking down *ATPα* in the eye also drove the expression of AMPs within the bodies of flies reared on either an AL or DR diet (**Fig. 3d**). Thus, DR fails to suppress immune responses in the context of forced photoreceptor degeneration.

Since we found that photoreceptor degeneration induced systemic immune responses, we postulated that reducing phototransduction should reduce inflammation. To assess how stress from environmental lighting influences immune responses, we analyzed a circadian microarray dataset comparing gene expression changes in wild-type (*y,w*) heads in flies reared in 12hr light and 12hr darkness (12:12LD) or constant darkness [32]. We found immune response genes to be among the most highly enriched processes upregulated in the flies housed in 12:12LD vs constant darkness (**Supplementary Fig. 3d, e and Supplemental Data 8**). We quantified AMPs within the bodies of flies harboring rhodopsin null mutations to evaluate how the different photoreceptor subtypes influence systemic immune responses. The *Drosophila* ommatidia consists of eight photoreceptors (R1-8) that express different rhodopsins with varying sensitives to distinct wavelengths of light [33]. The R1-6 photoreceptors express the major rhodopsin Rh1, encoded by *ninaE*, while the R7 photoreceptor expresses either Rh3 or Rh4. The R8 photoreceptor expresses either Rh5 or Rh6 [34]. The rhodopsin null mutants [*ninaE* [35]*, rh3* [36], *rh4* [37], or *rh6* [38]] displayed reductions in immune marker expression in their bodies compared to *w^1118^* outcrossed controls (**Fig. 3e**). Taken together, these findings indicate that suppression of rhodopsin mediated signaling is sufficient to suppress systemic immune responses in *Drosophila*.

Given the strong associations between chronic immune activation and accelerated aging, we examined the lifespans of nCLK-Δ flies [24]. Both nCLK-Δ1 and nCLK-Δ2 flies displayed significantly shortened lifespans, with a proportionally greater loss in median lifespan in flies reared on DR compared to AL (**Fig. 3f and Supplementary Fig. 3f-h**). nCLK-Δ flies have altered CLK function throughout all neurons, however, it is possible that the lifespan-shortening effect observed in these lines was substantially driven by loss of CLK-function within the eye; Others have demonstrated that CLK is highly enriched (>5-fold) within photoreceptors compared to other neuronal cell types in *Drosophila* (**Supplementary Fig. 3i**) [5]. Furthermore, over-expressing CLK-Δ1 within just photoreceptors (prCLK-Δ1) also shortened lifespan (**Supplementary Fig. 3j**). These findings argue that neuronal CLK function is required for the full lifespan extension mediated by DR and indicate that photoreceptor clocks are essential for maintenance of visual function with age and organismal survival.

### DR protects against lifespan shortening from photoreceptor cell stress

Previous reports have demonstrated that exposure to light can decrease lifespan— extending the daily photoperiod, or housing flies in blue light reduces longevity [39, 40]. Since DR delays visual senescence and promotes the rhythmic expression of genes involved in photoreceptor homeostasis (i.e., light adaptation, calcium handling), we investigated how diet influences survival in the context of light and/or phototransduction. To test the interrelationship among diet, light, and survival, we housed *w^1118^* (white-eyed) in either a 12:12 LD cycle or constant darkness. Housing flies in constant darkness extended the lifespan of flies reared on AL, while the lifespans of flies reared on DR were unaffected (**Fig. 4a**). Constant darkness failed to extend the lifespan of red-eyed (*w^+^*) *Canton-S* wild-type flies (**Supplementary Fig. 4a**), suggesting that the ATP-binding cassette transporter encoded by *w,* and the red-pigment within the cone-cells, helps to protect against lifespan shortening from diet- and light-mediated stress [41]. White-eyed, photoreceptor null flies (homozygous for *TRP^P365^* mutation [42]) reared on AL failed to display lifespan extension in constant darkness (**Supplementary Fig. 4b**), indicating that the lifespan shortening effects of light exposure are primarily mitigated by the photoreceptors.

**Figure 4.**
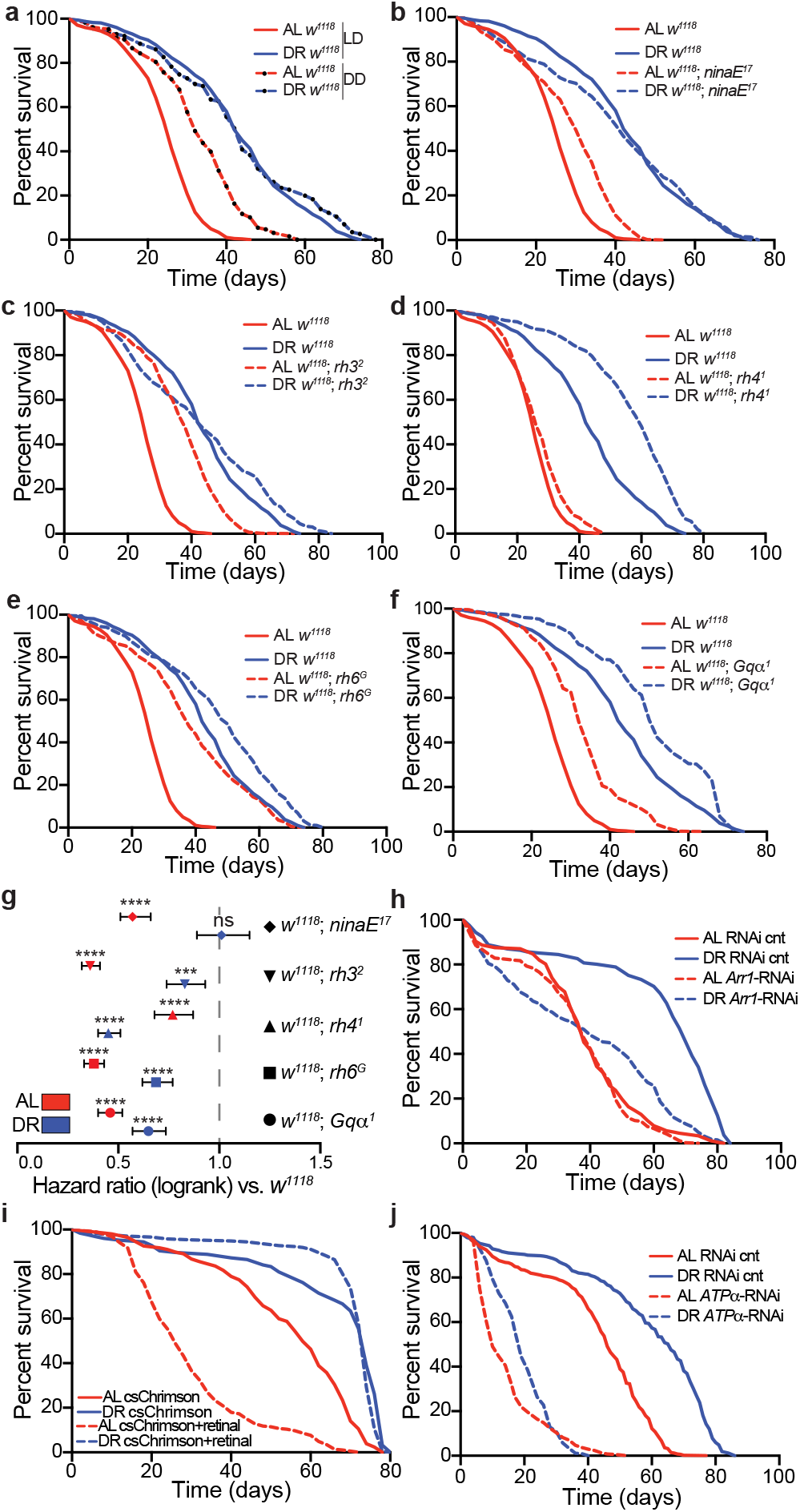
Photoreceptor activation modulates lifespan in a diet-dependent fashion. (**a**) Survival analysis of *w^1118^* flies housed in 12:12h LD or constant darkness (DD). Survival data is plotted as an average of three independent lifespan repeats. LD housed flies: AL *N*=560, DR *N*=584; DD: AL *N*=460, DR *N*=462. (**b**-**f**) Survival analysis of *w^1118^*; *ninaE^17^*, *w^1118^*; *rh3^2^*, *w^1118^*; *rh4^1^*, *w^1118^*; *rh6^G^*, and *w^1118^*; *Gq*α^1^ mutants compared to *w^1118^* control flies housed in 12:12h LD. Survival data is plotted as an average of three independent lifespan repeats. *Survival curves for *w^1118^* are re-plotted (b-f) for visual comparison, and the *w^1118^* and rhodopsin null lifespans repeats were performed simultaneously. All mutant lines were outcrossed to *w^1118^*. *w^1118^*; *ninaE^17^* flies: AL *N*=514, DR *N*=511; *w^1118^*; *rh3^2^* flies: AL *N*=543, DR *N*=597; *w^1118^*; *rh4^1^* flies: AL *N*=550, DR *N*=593; *w^1118^*; *rh6^G^* flies: AL *N*=533, DR *N*=563; *w^1118^*; *Gq*α*^1^* flies: AL *N*=403, DR *N*=400. (**g**) Hazard ratios for rhodopsin and Gq mutant flies compared to *w^1118^* control flies (ratios<1 indicate flies that are more likely to survive compared to *w^1118^*). Error bars indicate the 95% confidence interval of the hazard ratios. (**h**) Survival analysis of eye-specific *arr1*-RNAi knockdown flies vs RNAi control flies. Survival data is plotted as an average of two independent lifespan repeats for *arr1*-RNAi and one independent lifespan replicate for RNAi-controls. RNAi control flies: AL *N*=177, DR *N*=161; *arr1*-RNAi flies: AL *N*=333, DR *N*=322. (**i**) Survival analysis of retinal inducible, photoreceptor-specific optogenetic flies (Trpl-GAL4>UAS-csChrimson[red-shifted]) supplemented with retinal or vehicle control and housed in 12:12h red-light:dark. Survival data is plotted as an average of two independent lifespan repeats. Retinal treated flies: AL *N*=289, DR *N*=236; Vehicle treated flies: AL *N*= 256, DR *N*=126. (**j**) Survival analysis of eye-specific *ATP*α RNAi knockdown flies vs RNAi control flies. Survival data is plotted as an average of three independent lifespan repeats. RNAi control flies: AL *N*=493, DR *N*= 490; *ATP*α RNAi flies: AL *N*=510, DR *N*=535. (g) *P*values were determined by Log-rank (Mantel-Cox) test, ns denotes a non-significant *p*values, **** indicates *p*values less than 0.0001.

We performed survival analyses in rhodopsin null flies to examine how activation of the different photoreceptor subtypes influence lifespan on AL and DR. In agreement with the reduction in systemic immune responses observed in the rhodopsin null strains, these flies were also longer lived in comparison to *w^1118^* outcrossed controls (**Fig. 4b-e**). Furthermore, *rh6^G^* mutants, which displayed the largest reductions in inflammation, also displayed the greatest extension in lifespan compared to the other rhodopsin null lines. *Gqα^1^* mutants [43], which harbor a mutation in the G-protein that mediates activation of TRP channels downstream of rhodopsin, also displayed increased longevity compared to control flies (**Fig. 4f**). Interestingly, with the exception of Rh4, rhodopsin null mutations and *Gqα^1^* mutants primarily extended lifespan on AL, indicated by the hazard ratios in **Fig. 4g**. We next sought to investigate how increases in rhodopsin-mediated signaling influence survival. To this end, we knocked down the major arrestin protein, *arr1*, within the eyes of flies (GMR-GAL4>UAS-*arr1*-RNAi). Arr1 is required for light-mediated rhodopsin internalization from the rhabdomere membrane into endocytic vesicles, thus suppressing rhodopsin-mediated signaling and associated Ca^2+^-mediated phototoxicity/cell death [19, 44, 45]. In agreement with its physiological role in light-adaptation, we found that *arr1*-RNAi knockdown flies were hypersensitized to light (**Supplementary Fig. 4i**). In contrast to the Rhodopsin null strains which displayed greater proportional improvements in survival on AL vs DR, *arr1*-RNAi knockdown flies displayed significantly lifespan shortening on DR, while the lifespan on AL was indistinguishable from the control (**Fig. 4h**). Together, these data argue that DR-protects against lifespan shortening downstream of light and/or rhodopsin-mediated signaling in a manner that requires light-adaptation, and by extension, *arr1*-mediated rhodopsin endocytosis.

We utilized an optogenetics approach to examine how chronic photoreceptor activation influences survival in flies reared on AL or DR. Optogenetics is a powerful tool for examining how photoreceptor activation/suppression influences lifespan as it allows for the ability to compare lifespans within flies reared under the same lighting conditions, thus diminishing potential confounding variables present when comparing lifespan in different lighting conditions (i.e., LD vs constant darkness), such as extra-ocular effects of light on survival. To generate optogenetic flies we expressed the red-light-sensitive csChrimson cation channel [46] within photoreceptors (Trpl-GAL4>UAS-csChrimson). To activate the csChrimson channels, we housed the optogenetic flies in a 12:12 red-light:dark cycle and supplemented their food with either all-*trans* retinal (a chromophore required for full activation of csChrimson channels [47]) or a vehicle control (**Supplementary Fig. 4c**). Optogenetic activation of the photoreceptors (retinal treated) drastically reduced AL lifespan compared to vehicle treated controls, while the lifespan on DR was unaffected (**Fig. 4i**). Retinal did not appear to be toxic to flies lacking csChrimson channels, as the lifespan of *Canton-S* wild-type flies were indistinguishable between vehicle and retinal treated groups (**Supplementary Fig. 4d**).

Although DR protected flies from lifespan shortening from the optogenetic activation of photoreceptors, we found that forcing photoreceptor degeneration, by knocking down *ATPα* in the eye shortened lifespan on both AL and DR (**Fig. 4j**). Similarly, eye-specific knockdown of *nervana-2* and *-3* (*nrv2,* GMR-GAL4>UAS-*nrv2*-RNAi and *nrv3*, GMR-GAL4>UAS-*nrv3*-RNAi), which encode the *Beta* subunit of the Na^+^K^+^ATPase of the eye [31] also reduced phototaxis responses and shortened lifespan (**Supplementary Fig. 4e-h**). Taken together, these data support a model where DR protects flies from lifespan shortening caused by photoreceptor stress, as chronic photoreceptor activation reduces survival in flies reared on AL while having minimal to no effect on flies reared on DR. Inversely, photoreceptor deactivation primarily improves survival of flies reared on AL. ***Eye-specific, CLK-output genes modulate lifespan***

We next sought to determine if CLK-output genes in the eye influence age-related visual declines and lifespan. We employed a bioinformatics approach to identify candidate eye-specific circadian genes transcriptionally regulated by CLK (**Supplementary Fig. 5a and Supplemental Data 6**). First, we compared age-associated changes in photoreceptor-enriched gene expression [5] to genes that were differentially expressed on DR compared to AL. More than half of the photoreceptor-enriched genes that were downregulated with age were also upregulated on DR at ZT 0 and ZT 12 (**upper left quadrant of Fig. 5a, b and Supplementary Fig. 5a**). We then subset this gene list, selecting just transcripts whose expression was downregulated with age and upregulated on DR, and examined how their expression changed in nCLK-Δ1 fly heads (**Supplementary Fig. 5a**). From this analysis, we identified *Gβ76c*, *retinin,* and *sunglasses* as genes that were significantly downregulated in nCLK-Δ1 fly heads at ZT 0 and/or ZT 12 (**Fig. 5c, d and Supplementary Fig. 5a, e-g**). *Gβ76c* encodes the eye-specific G beta subunit that plays an essential role in terminating phototransduction [13, 48]. *Retinin* encodes one of the four most highly expressed proteins in the lens of the *Drosophila* compound eye [49]. Furthermore, *retinin* functions in the formation of corneal nanocoatings, knockdown of which results in degraded nanostructures and a reduction in their anti-reflective properties [50]. *Sunglasses*, also called *Tsp42Ej*, encodes for a lysosomal tetraspanin concentrated in the retina that protects against photoreceptor degeneration by degrading rhodopsin in response to light [51]. We analyzed a published CLK Chromatin Immunoprecipitation (ChIP-chip) dataset in flies and observed rhythmic CLK binding at the 5’-untranslated region of *sunglasses* in *Drosophila* eye tissue [52] (**Supplementary Fig. 5h and Supplementary Table 1**), which supports our bioinformatics approach and provides further evidence that *sunglasses* is an eye-specific CLK-output gene. Eye-specific knockdown of *Gβ76c* (GMR-GAL4>UAS-*Gβ76c*-RNAi)*, retinin* (GMR-GAL4>UAS-*retinin*-RNAi), and *sunglasses* (GMR-GAL4>UAS-*sunglasses*-RNAi) reduced phototaxis responses (**Fig. 5e and Supplementary Fig. 5i**), and shortened lifespan in comparison to RNAi control flies (GMR-GAL4>UAS-*mCherry*-RNAi) (**Fig. 5f, Supplementary Fig. 5j**). These findings indicate that DR and CLK function together in the regulation of eye-specific circadian genes involved in the negative regulation of rhodopsin signaling (i.e., phototransduction termination). Furthermore, these observations support previous findings that lifespan extension upon DR requires functional circadian clocks [10, 53], and establishes CLK-output genes as diet-dependent regulators of eye aging and lifespan in *Drosophila*.

**Figure 5.**
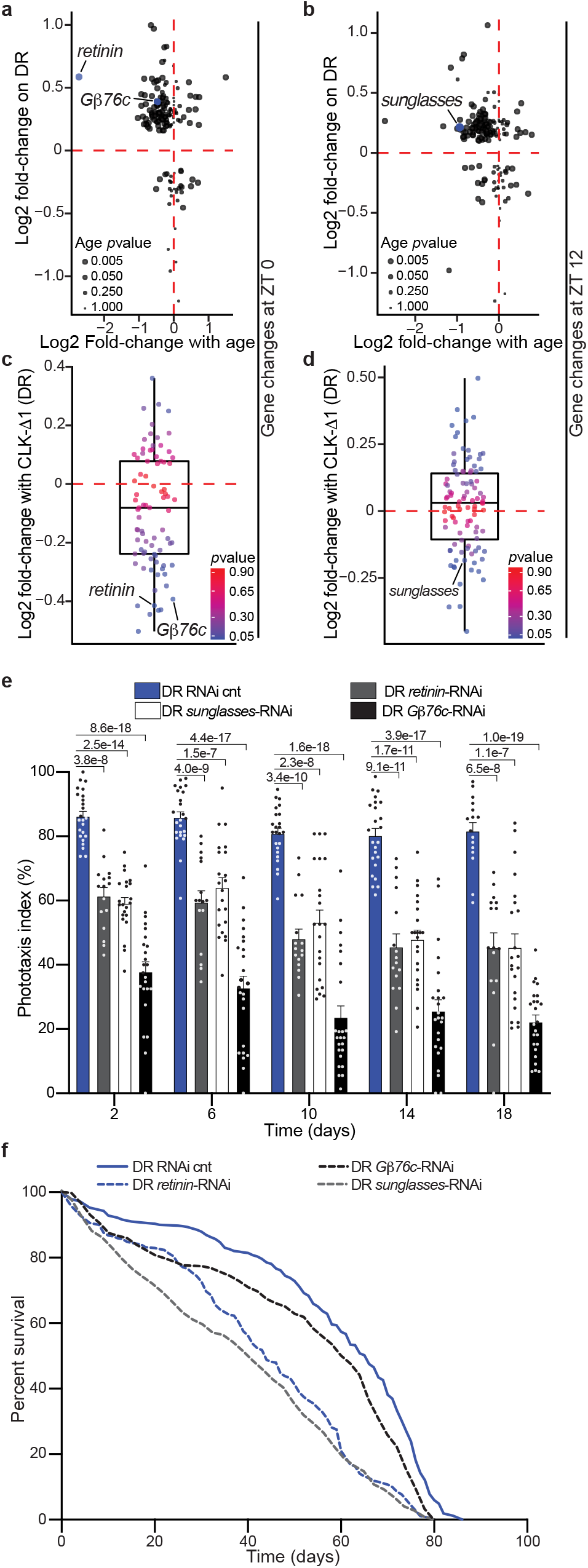
Knockdown of DR-sensitive, eye-specific CLK-output genes reduces survival. (**a**-**b**) Scatterplot of circadian, photoreceptor-enriched gene changes with age in wild-type heads (*x*-axis: 5- *vs* 55-day old flies) *vs* diet-dependent gene expression changes in heads from nCLK-Δ1 RNA-Seq control flies (*y*-axis: DR- *vs* AL-minus RU486) at ZT 0 (a) and ZT 12 (b). (**c**-**d**) Boxplots of the expression changes in nCLK-Δ1 heads (DR plus- vs DR minus-RU486) at ZT 0 (c) and ZT 12 (d) of genes that were downregulated with age and upregulated on DR (upper left quadrants of Fig. 5a, b). (**e**) Positive phototaxis responses with eye-specific knockdown of *G*β*76c* (GMR-GAL4> UAS-*G*β*76c*-RNAi), *retinin* (GMR-GAL4>UAS-*retinin*-RNAi), and *sunglasses* (GMR-GAL4>UAS-*sunglasses*-RNAi) compared to RNAi control flies (GMR-GAL4>UAS-*mCherry*-RNAi) reared on DR. For each timepoint results are represented as average phototaxis response +/-SEM (RNAi control *n*=24 biological replicates, *N*=480 flies per condition; *G*β*76c* RNAi *n*=24 biological replicates, *N*=480 flies per condition; *retinin* RNAi *n*=16 biological replicates, *N*=384 flies per condition; *sunglasses* RNAi *n*=24 biological replicates, *N*=480 flies per condition). (**f**) Survival analysis of eye-specific *G*β*76c*, *retinin*, *sunglasses*, and RNAi knockdown flies compared to RNAi control flies reared on DR. Survival data is plotted as an average of three independent lifespan repeats for RNAi control, *sunglasses*, and *G*β*76c* flies and two independent lifespan repeats for *retinin* RNAi knockdown flies. RNAi-cnt flies: *N*=490; *retinin*-RNAi flies: *N*=363; *sunglasses*-RNAi flies: *N*=468; *G*β*76c*-RNAi flies: *N*=509. (e) *P*values were determined by two-tailed Student’s ttest (unpaired) at each timepoint comparing the phototaxis index of RNAi control flies to *G*β*76c*-, *retinin*-, and *sunglasses*-RNAi flies.

## Discussion

Progressive declines in circadian rhythms are one of the most common hallmarks of aging observed across most lifeforms [54]. Quantifying the strength, or amplitude, of circadian rhythms is an accurate metric for predicting chronological age [55]. Many cellular processes involved in aging (e.g., metabolism, cellular proliferation, DNA repair mechanisms, etc.) display robust cyclic activities. Both genetic and environmental disruptions to circadian rhythms are associated with accelerated aging and reduced longevity [56, 57]. These observations suggest that circadian rhythms may not merely be a biomarker of aging; rather, declines in circadian rhythms might play a causal role. The observation that DR and DR-memetics, such as calorie restriction and time-restricted feeding, improve biological rhythms suggests that clocks may play a fundamental role in mediating their lifespan-extending benefits.

Herein, we identified circadian processes that are selectively amplified by DR. Our findings demonstrate that DR amplifies circadian homeostatic processes in the eye, some of which are required for DR to delay visual senescence and improve longevity in *Drosophila*. Taken together, our data demonstrate that photoreceptor stress has deleterious effects on organismal health; disrupting CLK function and/or overstimulation of the photoreceptors induced a systemic immune response and reduced longevity. Our findings establish the eye as a diet-sensitive regulator of lifespan. DR’s neuroprotective role in the photoreceptors appears to be mediated via the molecular clock, which promotes the rhythmic oscillation of genes involved in the suppression of phototoxic cell stress (**Fig. 6 and Supplemental Discussion 1**). Our data also support the idea that age-related declines in the visual system impose a high cost on an organism. Perhaps this is why a number of long-lived animals have visual systems that have undergone regressive evolution (e.g., cave-dwelling fish and naked-mole rats) [58]. Failing to develop a visual system may help these organisms avoid age-related damage and inflammation caused by retinal degeneration. Ultimately, developing a visual system, which is critical for reproduction and survival, may be detrimental to an organism later in life. Thus, vision may be an example of an antagonistically pleiotropic mechanism that shapes lifespan.

**Fig. 6.**
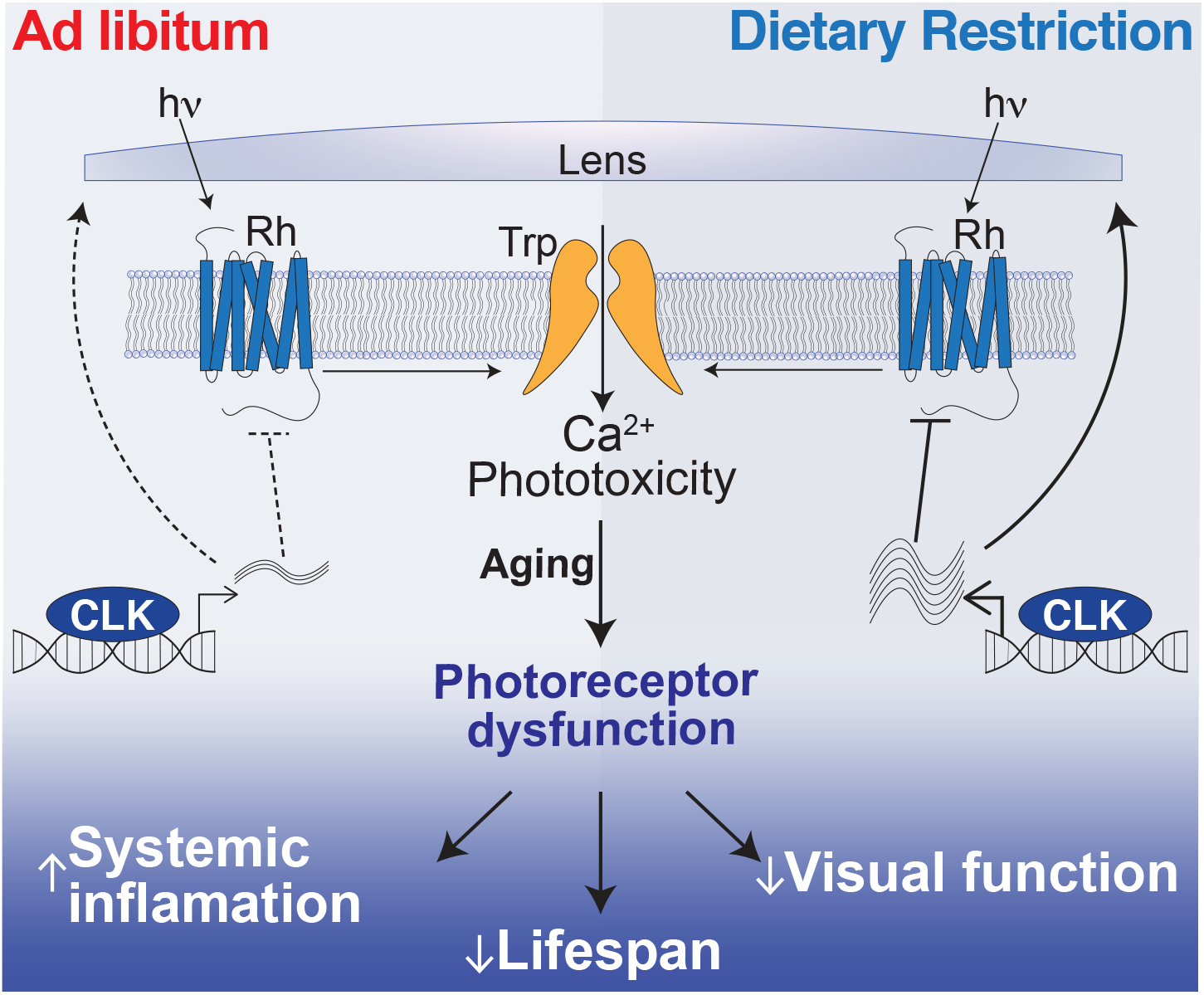
Dietary restriction extends lifespan by promoting rhythmic homeostatic processes in the eye. DR promotes CLK-output processes in the eye that suppress light/Ca^2+^-mediated phototoxicity to suppress systemic inflammaion, delay visual senesence, and improve survival.

## Methods

### Fly stocks

The genotypes of the *Drosophila* lines used in this study are listed in Supplemental Table 2. The following lines were obtained from the Bloomington *Drosophila* Stock Center: *Oregon R.* (25125), GMR-GAL4 (1104), Elav-GS-GAL4 (43642), Trpl-GAL4 (52274), *Clk^OUT^* (56754), UAS-csChrimson (55134), UAS-CLK-Δ1 (36318), UAS-CLK-Δ2 (36319), *Gβ76c*-RNAi (28507), *tsp42Ej/sunglasses*-RNAi (29392), *retinin*-RNAi (57389), *ATPα*-RNAi (28073), *nrv2*-RNAi (28666), *nrv3*-RNAi (60367), and mCherry-RNAi (Bloomington RNAi-cnt, 35785). The following lines were obtained from the Vienna *Drosophila* Resource Center: *arr1*-RNAi (22196), *RNAi*-cnt (empty vector, 60100). The following lines were received from the laboratory of Craig Montell: *CantonS*, *w^1118^*, *w^1118^*; *ninaE^17^*, *w^1118^*; *rh3^2^*, *w^1118^*; *rh4^1^*, *w^1118^*; *rh6^G^*, *w^1118^*; *Gqα^1^*, and TRP^365^. The following lines were outcrossed to *w^1118^* for this manuscript: UAS-CLK-Δ1^OC^ and *CantonS*^OC^. The Trpl-GAL4 line was recombined with GAL80 for this manuscript: Trpl-GAL4; GAL80^ts^.

### Fly husbandry and survival analyses

All flies were maintained at 25±1 °C, 60% humidity under a 12h:12h LD cycle (∼750lux, as measured with a Digital Lux Meter, Dr. Meter Model LX1330B) unless otherwise indicated. Fly stocks and crosses were maintained on a standard fly media as described previously [59]. Briefly, standard fly media consisted of 1.5% yeast extract, 5% sucrose, 0.46% agar, 8.5% of corn meal, and 1% acid mix (a 1:1 mix of 10% propionic acid and 83.6% phosphoric acid). Fly bottles were seeded with live yeast prior to collecting virgins or setting up crosses. Mated adult progeny were then transferred to *ad libitum* (AL) or dietary restriction (DR) media within three days of eclosion. Adult flies used in experiments were transferred to fresh media every 48h at which point deaths were recorded for survival analysis. AL and DR fly media differed only in its percentage of yeast extract, respectively containing 5% or 0.5% (Yeast Extract, B.D. Bacto, Thermo Scientific 212720, Cat no. 90000-722). *Optogenetic experiments*: For experiments using the csChrimson channel rhodopsin [46], adult flies were transferred to media supplemented with 50μM all-trans-retinal (Sigma Aldrich, R2500-1G) or drug vehicle (100% ethanol), and maintained under a 12h:12h red light:dark cycle, with ∼10lux of red light (∼590nm) during the light phase. *Elav-GeneSwitch flies*: GeneSwitch [60], adult flies were transferred to media supplemented with 200μM RU486 (Mifepristone, United States Biological), indicated as either AL+ or DR+, for post-developmental induction of transgenic elements; isogenic control flies were transferred to food supplemented with a corresponding concentration of drug vehicle (100% ethanol), indicated as either AL- or DR-. *prCLK-Δ1 experiments*: GAL80 temperature sensitive crosses were set in bottles at 25°C, 60% humidity under a 12h:12h LD cycle for four days. Parental flies were removed, and the bottles were transferred to 18°C for approximately three-weeks to suppress GAL4 activity throughout development. After ecolsion, the F1 generations were sorted onto AL or DR food the flies were maintained at 30°C to de-repress GAL80 and activate GAL4 (60% humidity under a 12h:12h LD cycle) for the remainder of their lifespans. The F1 generations for these experiments share the same genetic background, as both the UAS-CLK-Δ1 and the *CantonS* control lines were fully outcrossed to the same *w^1118^* strain prior to setting up the cross with Trpl-GAL4; GAL80^ts^.

### Circadian time-course expression analysis

Mated *Canton-S* females were reared on AL or DR diets for seven days at 25±1 °C, under a 12:12h light-dark (LD) regimen. Beginning on the seventh day, four independent biological replicates (per diet/timepoint) of approximately 35 flies were collected on dry ice every four-hours for 20-hours starting at ZT 0 (six total timepoints, 48 total samples). RNA extraction, DNA amplification/labeling, and gene expression arrays were performed following the same protocols as in Katewa *et al.*, 2012 [61]. In summary, RNA was isolated from whole fly lysates with Qiagen’s RNeasey Lipid Tissue Mini Kit (74804) and RNA quantity and quality were accessed with a Nanodrop and Agilent’s bioanalyzer (RNA 600 Nano Kit (5067-15811)). DNA amplification from total RNA was performed using Sigma’s TransPlex Complete Whole Transcriptome Amplification Kit (WTA2) and purified with Qiagen’s QIAquick PCR Purification Kit (28104). Gene expression labeling was performed with NimbleGen One-Color DNA Labeling Kit (05223555001) and hybridized to NimbleGen 12-Plex gene expression arrays. Arrays were quantitated with NimbleGen’s NimbleScan2 software, and downstream expression analyses were conducted in R (http://www.r-project.org). Transcript-level expression from the four independent biological replicates were averaged for each time-point.

### nCLK-Δ1 RNA-Seq analyses

nCLK-Δ1 (Elav-GS-GAL4>UAS-nCLK-Δ1) adult flies were developed on standard stock food (1.5% yeast-extract) for four days. Three independent biological replicates of 100 mated female flies were then reared on AL or DR diets treated with RU486 or vehicle control at 25±1 °C, under a 12:12h LD regimen. Diets were changed approximately every 48-hours, until the seventh day at which point flies were flash frozen on dry-ice at ZT 0 and ZT 12 (lights-on and -off, respectively). See Supplemental Fig. 2a for RNA-Seq. experimental design. *RNA-extraction*: Frozen flies were vortexed to remove heads and mRNA from each biological replicate of pooled heads was isolated with the Quick-RNA MiniPrep Kit (Zymo Research #11-328), per manufactures’ instructions. *Fragment library preparation and deep sequencing*: Library preparation was performed by the Functional Genomics Laboratory (FGL), a QB3-Berkeley Core Research Facility at University of California, Berkeley. cDNA libraries were produced from the low-input RNA using the Takara SMART-Seq v4 Ultra-low input RNA kit. An S220 Focused-Ultrasonicator (Covaris®) was used to fragment the DNA, and library preparation was performed using the KAPA hyper prep kit for DNA (KK8504). Truncated universal stub adapters were used for ligation, and indexed primers were used during PCR amplification to complete the adapters and to enrich the libraries for adapter-ligated fragments. Samples were checked for quality on an AATI (now Agilent) Fragment Analyzer. Samples were then transferred to the Vincent J. Coates Genomics Sequencing Laboratory (GSL), another QB3-Berkeley Core Research Facility at UC Berkeley, where Illumina sequencing library molarity was measured with quantitative PCR with the Kapa Biosystems Illumina Quant qPCR Kits on a BioRad CFX Connect thermal cycler. Libraries were then pooled evenly by molarity and sequenced on an Illumina NovaSeq6000 150PE S4 flowcell, generating 25M read pairs per sample. Raw sequencing data was converted into fastq format, sample specific files using the Illumina bcl2fastq2 software on the sequencing centers local linux server system. *Read alignment and differential expression analyses*: Raw fastq reads were filtered by the Trimmomatic software [62] (Trimmomatic-0.36) to remove Illumina-specific adapter sequences and the minimal length was set to 36 (MINLEN) for trimming sequences. The paired end filtered reads were then aligned to the *D. Melanogaster* dm6 genome (BDGP Release 6 + ISO1 MT/dm6) by HISAT2 [63] to generate BAM files with the specific strand information set to “Reverse”. Count files were then generated by featureCounts [64] and the *D. Melanogaster* reference genome was utilized as the gene annotation file with specific strand information set to “stranded (Reverse)”. Resulting count files (tabular format) were then analyzed with DEseq2 [65] with fit-type set to “local”, and *p*values of less than 0.05 were considered differentially expressed between factor levels. Normalized count reads were outputted for visualization of expression (heatmaps), and Supplemental Data Files 3a contains normalized count reads across all experimental samples. *UCSC genome browser visualization*: The makeUCSCfile software package from HOMER was utilized to generate bedGraph files for visualizing changes in tag density at exon 2 of *clk* comparing nCLK-Δ1 and control samples (Supplementary Fig. 2B).

### Heatmap visualizations

We employed the heatmap2 function from R gplots package to visualize bioinformatics data. Data were not clustered, and data were scaled by row for normalization across time-points.

### Electroretinogram assays

ERGs were performed and analyzed in two independent laboratories. ERGs were recorded for nCLK-Δ1 flies reared on AL or DR diets supplemented with vehicle or RU486 at day 14 at the Baylor College of Medicine (BCM), and at day 21 at the University of California, Santa Barbara (UCSB). ERGs were recorded for prCLK-Δ1 flies at UCSB reared on AL or DR and maintained at either 18°C or 30°C. *BCM*: ERG recordings were performed as in Wang *et al*., 2014 [66]. Flies were glued on a glass slide. A recording electrode was placed on the eye and a reference electrode was inserted into the back of the fly head. Electrodes were filled with 0.1 M NaCl. During the recording, a 1 s pulse of light stimulation was given. The ERG traces of at least eight flies per genotype/diet were recorded and analyzed by LabChart8 software (AD Instruments). *UCSB*: ERG recordings were performed as in Wes *et al*., 1999 [67]. Two glass electrodes were filled with Ringer’s solution and electrode cream was applied to immobilized flies. A reference electrode was placed on the thorax, while the recording electrode was placed on the eyes. Flies were then exposed to a 10s pulse of ∼200lux white light, a light intensity that is comparable to the phototaxis assay. An EI-210 amplifier (Warner Instruments) was used for amplifying the electrical signal from the eye after light stimulation, and the data were recorded using a Powerlab 4/30 device along with the LabChart 6 software (AD Instruments). Raw data were then uploaded into R-statistical software for plotting and statistical analysis. All electroretinograms were performed between ZT4-8 or ZT12-14.

### Positive phototaxis assay

Positive phototaxis was performed using an adapted protocol from Vang *et al.*, 2014 [68] (Fig S2D). Phototaxis measurements were recorded longitudinally on populations of female flies aged on either AL or DR food (with or without 200μM RU486 when indicated) at a density of 10-25 flies per tube prior to and after phototaxis measurements. On the day of phototaxis recording, eight groups of flies (four AL and four DR groups) were placed in separate 2.5cm x 20cm tubes (created from three enjoined narrow fly vials [Genesee Scientific]) and dark-adapted for 15-minutes prior to light exposure (no food was available in the vials during phototaxis assays). Flies were then gently tapped to the bottom of the tube, placed horizontally, and exposed to white light from an LED strip (Ustellar, UT33301-DW-NF). A gradient of light intensity was created, with 500lux at the nearest point in the fly tube to the light source and 150lux at the furthest point. Phototaxis activity was recorded by video at 4K resolution (GoPro, Hero5 black). Positive phototaxis was scored manually as the percentage of flies that had traveled >19cm toward the light source in three 15s intervals (15s, 30s, and 45s). “Phototaxis index” was calculated by averaging the percent of positive phototaxis for each vial at the three 15s intervals. To control for light-independent wandering activity, a phototaxis index was also calculated when the light source was placed in parallel to the fly tube, such that all parts of the tube were equally illuminated with 500lux. *Normalizing phototaxis responses to wandering activity failed to significantly affect phototaxis index, data not shown.

### RNA extraction and cDNA preparation

Flies were maintained on AL or DR for the indicated amount of time, then flash frozen on dry ice. Heads were separated from bodies (thorax and abdomen) by vigorous shaking. Flies were then ground using a hand-held homogenizer at room temperature following MiniPrep instructions. Total RNA was isolated using the Quick-RNA MiniPrep Kit (Zymo Research, 11-328). In brief, flies were maintained on AL or DR for the indicated amount of time, then flash frozen on dry ice. Heads were separated from bodies (thorax and abdomen) by vigorous shaking. Flies were then ground using a hand-held homogenizer at room temperature following MiniPrep instructions. RNA was collected into 30μl DNAse/RNAse-free water and quantified using the NanoDrop 1000 Spectrophotometer (Thermo Scientific). For each experiment, 120-180 age-, genotype-, and diet-matched flies were collected, and three independent RNA extractions were performed. To extract RNA from heads, 40-60 flies were used; to extract RNA from bodies, 20-30 flies were used. *cDNA preparation*: The iScript Reverse Transcription Supermix for RT-qPCR (Bio-Rad, 1708841) was used to generate cDNA from RNA extracted from heads and bodies. For each group, 1 μg of total RNA was placed in a volume of 4μl iScript master mix, then brought to 20μl with DNAse/RNAse-free water. A T1000 thermocycler (BioRad) was used for first-strand RT-PCR reaction following iScript manufacturers’ instructions—priming step (5min at 25°C), reverse transcription (30min at 42°C), and inactivation of the reaction (5min at 85°C).

### Real-time quantitative PCR

Reactions were performed in a 384-well plate. Each reaction contained 2μl of 1:20 diluted cDNA, 1μl of primers (forward and reverse at 10μM), 5μl SensiFAST SYBR Green No-ROX Kit (BIOLINE, BIO-98020), and 2μl of DNAse/RNAse-free water. The qPCR reactions were performed with a Light Cycler 480 Real-Time PCR machine (Roche Applied Science) with the following run protocol: pre-incubation (95°C for 2 min), forty PCR cycles of denaturing (95°C for 5s, ramp rate 4.8°C/s), and annealing and extension (60°C for 20 s, ramp rate 2.5°C/s).

### Hemolymph Mass spectrometry

*Proteomic sample preparation*: nCLK-Δ1 female flies (Elav-GeneSwitch-GAL4>UAS-nCLK-Δ1) were reared on AL diet plus RU486 or vehicle control (*N*=300 flies per biological replicate, *n*=3 biological replicates). At day 14, flies were snap frozen on dry ice and transferred to pre-chilled vials. The vials were vortexed for 5-10s to remove heads and the frozen bodies were transferred to room temperature vials fitted with 40µm filters. Headless bodies were thawed at room temperature for 5 minutes and spun at 5000 rpm for 10min at 4 °C. Following the spin, hemolymph collected at the bottom of each vial and the bodies remained within the filters. *Digestion*: A Bicinchoninic Acid protein assay (BCA) was performed for each of the hemolymph samples and a 100µg aliquot was used for tryptic digestion for each of the 6 samples. Protein samples were added to a lysis buffer containing a final concentration of 5% SDS and 50 mM triethylammonium bicarbonate (TEAB), pH ∼7.55. The samples were reduced in 20 mM dithiothreitol (DTT) for 10 minutes at 50⁰ C, subsequently cooled at room temperature for 10 minutes, and then alkylated with 40 mM iodoacetamide (IAA) for 30 minutes at room temperature in the dark. Samples were acidified with a final concentration of 1.2% phosphoric acid, resulting in a visible protein colloid. 90% methanol in 100 mM TEAB was added at a volume of 7 times the acidified lysate volume. Samples were vortexed until the protein colloid was thoroughly dissolved in the 90% methanol. The entire volume of the samples was spun through the micro S-Trap columns (Protifi) in a flow-through Eppendorf tube. Samples were spun through in 200 µL aliquots for 20 seconds at 4,000 x g. Subsequently, the S-Trap columns were washed with 200 µL of 90% methanol in 100 mM TEAB (pH ∼7.1) twice for 20 seconds each at 4,000 x g. S-Trap columns were placed in a clean elution tube and incubated for 1 hour at 47⁰ C with 125 µL of trypsin digestion buffer (50 mM TEAB, pH ∼8) at a 1:25 ratio (protease:protein, wt:wt). The same mixture of trypsin digestion buffer was added again for an overnight incubation at 37⁰ C. Peptides were eluted from the S-Trap column the following morning in the same elution tube as follows: 80 µL of 50 mM TEAB was spun through for 1 minute at 1,000 x g. 80 µL of 0.5% formic acid was spun through next for 1 minute at 1,000 x g. Finally, 80 µL of 50% acetonitrile in 0.5% formic acid was spun through the S-Trap column for 1 minute at 4,000 x g. These pooled elution solutions were dried in a speed vac and then re-suspended in 0.2% formic acid. *Desalting*: The re-suspended peptide samples were desalted with stage tips containing a C18 disk, concentrated and re-suspended in aqueous 0.2% formic acid containing “Hyper Reaction Monitoring” indexed retention time peptide standards (iRT, Biognosys). *Mass spectrometry system*: Briefly, samples were analyzed by reverse-phase HPLC-ESI-MS/MS using an Eksigent Ultra Plus nano-LC 2D HPLC system (Dublin, CA) with a cHiPLC system (Eksigent) which was directly connected to a quadrupole time-of-flight (QqTOF) TripleTOF 6600 mass spectrometer (SCIEX, Concord, CAN). After injection, peptide mixtures were loaded onto a C18 pre-column chip (200 µm x 0.4 mm ChromXP C18-CL chip, 3 µm, 120 Å, SCIEX) and washed at 2 µl/min for 10 min with the loading solvent (H_2_O/0.1% formic acid) for desalting. Subsequently, peptides were transferred to the 75 µm x 15 cm ChromXP C18-CL chip, 3 µm, 120 Å, (SCIEX), and eluted at a flow rate of 300 nL/min with a 3 h gradient using aqueous and acetonitrile solvent buffers. *Data-dependent acquisitions (for spectral library building)*: For peptide and protein identifications the mass spectrometer was operated in data-dependent acquisition [51] mode, where the 30 most abundant precursor ions from the survey MS1 scan (250 msec) were isolated at 1 m/z resolution for collision induced dissociation tandem mass spectrometry (CID-MS/MS, 100 msec per MS/MS, ‘high sensitivity’ product ion scan mode) using the Analyst 1.7 (build 96) software with a total cycle time of 3.3 sec as previously described [69]. *Data-independent acquisitions*: For quantification, all peptide samples were analyzed by data-independent acquisition (DIA, e.g. SWATH), using 64 variable-width isolation windows [70, 71]. The variable window width is adjusted according to the complexity of the typical MS1 ion current observed within a certain m/z range using a DIA ‘variable window method’ algorithm (more narrow windows were chosen in ‘busy’ m/z ranges, wide windows in m/z ranges with few eluting precursor ions). DIA acquisitions produce complex MS/MS spectra, which are a composite of all the analytes within each selected Q1 m/z window. The DIA cycle time of 3.2 sec included a 250 msec precursor ion scan followed by 45 msec accumulation time for each of the 64 variable SWATH segments.

### Identification of photoreceptor enriched CLK-output genes

Diagram of bioinformatics steps reported in Supplementary Fig. 5A. Gene-lists are reported in **Supplemental Data 6**. We identified the top 1,000 photoreceptor-enriched genes from Charlton-Perkins *et al*., 2017 [72] (GSE93782). We then filtered this list for genes that oscillate in a circadian fashion, and that are downregulated with age from Kuintzle *et al*., 2017 [15] (GSE81100). Approximately 1/3 of the photoreceptor enriched genes (366 genes) were expressed in a circadian fashion in young wild-type heads and approximately one-half of these (172 genes) displayed a significant loss in expression with age (5- vs 55-day old heads). We further analyzed the remaining gene lists to identify those that are significantly upregulated on DR compared to AL at either ZT 0 or ZT 12 from control (vehicle treated) samples from our nCLK-Δ1 RNA-Seq analyses. For the final filtering step, we analyzed the genes that were significantly downregulated in nCLK-Δ1 on DR (RU486 vs vehicle treated controls), resulting in the identification of *Gβ76c*, *retinin*, and *sunglasses*.

### Statistical analysis

The individual biological replicates “*n*” and the number of individual flies *“N*” is denoted in each figure legend along with the particular statistical test utilized. The *p*value statistics are included in each figure. All error bars are represented as standard error of the mean (SEM), and all graphs were generated in PRISM 9 (GraphPad). The experiments in this manuscript were performed with populations of female flies (i.e., typically greater than 20 flies per technical replicate).

#### Time-course microarray analyses

Four independent biological replicates (per diet/timepoint) of approximately 35 *CantonS* female flies were collected on dry ice every four-hours for 20-hours starting at ZT 0 (six total timepoints, 48 total samples). Differential expression was determined by two-tailed Student’s *t*test (paired) comparing the averaged transcript-level expression values between AL and DR samples across all timepoints, and *p*values less than 0.05 were considered significant. The JTK_CYCLE algorithm [73] (v3.0) was utilized to identify circadian transcripts from the AL and DR time-course expression arrays. Transcript level expression values for each of the four biological replicates (per timepoint/diet) were used as input for JTK_CYCLE, and period length was set to 24-hours. We defined circadian transcripts as those displaying a JTK_CYCLE *p*value of less than 0.05. Subsequent analyses compared diet-dependent changes in JTK_CYCLE outputs (phase and amplitude).

#### nCLK-Δ1 RNA-Seq

Three independent biological replicates of 100 mated female adult flies were utilized per genotype/diet/time-point. DEseq2 software [65] was utilized and *p*values of less than 0.05 were considered differentially expressed between factor levels.

#### ERG responses

For ERG experiments we quantified responses from 6-15 individual flies per standards in the field. Statistical significance was determined by two-tailed Student’s *t*test (unpaired), comparing ERG responses between diet and genotypes. Full ERG statistics are reported in **Supplemental Data 10**.

#### Survival analyses

The Log-rank (Mantel-Cox) test was used to determine statistical significance comparing average lifespan curves from a minimum of two independent lifespan replicates. Hazard Ratios (logrank) were also utilized to determine the probability of death across genotypes, lighting conditions, and diet. Detailed Log-rank and hazard ratios for each lifespan are reported in **Supplemental Data 7**.

#### Positive phototaxis assay

Statistical significance for phototaxis index at each timepoint were calculated with the Student’s *t*test (two-tailed, un-paired). 2way ANOVA or mixed-effects models were performed to determine statistical significance between diet, genotype, or time interactions. Full statistical output (2way ANOVA and *t*test) for all phototaxis experiments is reported in **Supplemental Data 9**.

#### Real time quantitative PCR

Fold-change in gene expression was calculated using the ΔΔCt method and the values were normalized using *rp49* as an internal control. *P*values were calculated with the pairwise Student’s *t*-test comparing Log2 fold-changes in expression.

#### Mass-spectrometric data processing, quantification and bioinformatics

Mas spectrometric data-dependent acquisitions [51] were analyzed using the database search engine ProteinPilot (SCIEX 5.0 revision 4769) using the Paragon algorithm (5.0.0.0,4767). Using these database search engine results a MS/MS spectral library was generated in Spectronaut 14.2.200619.47784 (Biognosys). The DIA/SWATH data was processed for relative quantification comparing peptide peak areas from various different time points during the cell cycle. For the DIA/SWATH MS2 data sets quantification was based on XICs of 6-10 MS/MS fragment ions, typically y- and b-ions, matching to specific peptides present in the spectral libraries. Peptides were identified at Q< 0.01%, significantly changed proteins were accepted at a 5% FDR (*q*-value < 0.01).

#### Gene-ontology enrichment analysis

To identify enriched gene-ontology (i.e., bioprocess) categories with the resultant lists from bioinformatics approaches, we utilized the “findGO.pl” package from HOMER. Full gene-ontology lists including enrichment statistics and associated gene-lists are reported in supplemental data files. A maximal limit of 200 gene identifiers per GO category was implemented to reduce the occurrence of large, over-represented terms that lack specificity (i.e., *metabolism*). Full gene-ontology lists are reported in supplemental data files.

### Data availability

Time-course microarray data and accompanied JTK_CYCLE statistics that support the findings in this study have been deposited in the Gene Expression Omnibus [74] with the GSE158286 accession code. The RNAseq data and accompanied differential expression analyses that support the findings in this study have been deposited to GEO with the GSE158905 accession code. The mass spectrometric raw data are deposited at ftp://MSV000086781@massive.ucsd.edu (MassIVE user ID: MSV000086781, password: winter; preferred engine: Firefox); it is also available at ProteomeXchange with the ID PXD023896. Additional mass spectrometric details from DIA and DDA acquisitions, such as protein identification and quantification details are available at the repositories (including all generated Spectronaut and Protein Pilot search engine files).

## Acknowledgements

We would like to thank the QB3 Genomics Functional Genomics Lab at UC Berkeley and the Vincent J. Coates Genomics Sequencing Lab for RNA-seq library preparation and deep sequencing. We thank the Bloomington *Drosophila* Stock Center, the Vienna *Drosophila* Resource Center, and Craig Montell for providing flies used in this study. We would like to acknowledge Hugo Bellen and Zhongyuan Zuo for contributing resources and insight. We thank Daron Yim and John McCarthy for their assistance. We would like to acknowledge the work of Paolo Sassone-Corsi and his recent contributions to the fields on nutrition, circadian rhythms, and aging that helped in the conception of this project. This work is funded by grants awarded to P.K. from the American Federation of Aging Research, NIH grants R01 R01AG038688 and AG045835 and the Larry L. Hillblom Foundation. B.A.H. is supported by NIH/NIA T32 award AG000266 and C.M. is supported by NIH/NEI awards EY008117 and EY010852. We acknowledge the Buck Institute Proteomics Core and the support of instrumentation from the NCRR shared instrumentation grant 1S10 OD016281.

## Author Contributions

Conceptualization, B.A.H., G.T.M., and S.K.; Software, B.A.H., G.T.M., B.S., and S.K.; Formal analysis, B.A.H., G.T.M., and B.S.; Investigation, B.A.H., G.T.M., C.L., T.L., N.L., D.L-K., and S.B.; Visualization, B.A.H.; Writing-Original Draft, B.A.H., Writing-Review & Editing, B.A.H., G.T.M., and P.K.; Data Curation, B.A.H., G.T.M., S.M., and B.S.; Methodology, G.T.M. and M.L., Resources, C.M. and P.K., Supervision, P.K., Funding acquisition, C.M. and P.K.

## Additional information

The authors declare no competing interests.

**Supplementary Information** is available for this paper Correspondence and requests for materials should be addressed to P.K.

## Supplemental Discussion 1

### DR amplifies and modifies circadian transcriptional output

DR and DR-memetics have long been known to improve circadian behavioral rhythms in old age [1]. Over the past decade, improvements in molecular genetic techniques and next-generation sequencing have allowed investigators to examine how nutrient composition and time of feeding influence circadian transcriptional rhythms. Reports in mammals have demonstrated that calorie restriction, a reduction in total calorie intake without malnutrition, enhances the number and amplitude of rhythmic transcripts [2]. Inversely, high-nutrient diets, such as high-fat/western diets suppress circadian transcriptional rhythms [3]. The near doubling in the number of circadian transcripts we quantified on DR vs AL in *Drosophila* is consistent with observations in mammals. Additionally, transcripts oscillating on DR displayed an increased circadian amplitude. DR-mediated increases in the number of circadian transcripts, and their amplitude, is likely due to enhanced transcriptional output by CLK/CYC. Recent reports in both mice and flies have demonstrated that nutrient-sensing mechanisms (i.e., AMPK/TOR and Sirtuin signaling) signal directly to the core-clock transcription factors to activate transcription [4–6]. For instance, the *Drosophila* AMPK, which is activated in response to cellular energy depletion (e.g., elevated AMP concentrations), directly phosphorylates CLK, enhancing its circadian transcriptional output [4]. Because we extracted mRNA from populations of flies, the transcript expression values we report here are influenced by both individual and population-wide transcript expression levels. Therefore, the DR-mediated improvements to the circadian transcriptome we have observed may also reflect greater synchronicity between individual flies.

Interestingly, we also observed relatively low overlap between the transcripts that oscillate on AL compared to DR. We found that DR-oscillating genes are enriched for processes related to homeostatic function, while circadian processes on AL-oscillating genes are enriched for processes related to damage-response pathways. These findings are similar to those reported in mouse liver tissue, comparing transcriptome of mice reared on standard chow versus those on calorie restriction [2]. A combination of aging and damage response signals (e.g., reactive oxygen species) also influence which transcripts cycle in the fly [7]. This phenomenon, termed “circadian reprogramming,” is also observed in response to nutrient cues, where differing nutrient signals direct which specific transcripts are transcriptionally targeted downstream of the molecular clock. The similarities between the diet-dependent changes we report here and those previously reported in mice on calorie restriction indicates that the molecular clock’s response to nutrient restriction is evolutionarily conserved.

Given DR’s ability to robustly extend lifespan while amplifying circadian transcriptional output, we postulated that DR-sensitive circadian processes play an important role in slowing aging and improving survival. Although highly informative, to date, the diet-dependent circadian transcriptome studies have analyzed only a small number of mammalian tissues and thus have provided only a limited description of how diet influences circadian transcriptional output at the whole-organism level. Our ability to analyze the AL/DR circadian transcriptomes in the whole fly allowed for an unbiased approach for identifying the most DR-sensitive, cyclic processes throughout the body. This approach led to the observation that phototransduction was among the top circadian processes amplified by DR. The phototransduction genes we identified were also cyclic in flies reared on AL, albeit at a lower expression and circadian amplitude, indicating that their transcriptional regulation is likely not a result of circadian reprogramming. This, however, highlights the biological importance of their circadian regulation. A limitation of our AL/DR circadian transcriptome analyses is that they are likely under-powered to identify the full spectrum of eye-specific circadian transcripts, because our mRNA samples were pooled from whole-body lysates and were collected for only one circadian cycle (24hr). Analyses of a more robust circadian transcriptome, performed from mRNA collected from heads, over 2 circadian cycles (48hr), indicated that phototransduction components were among the most rhythmic circadian processes, thus underscoring the importance of circadian regulation within the eye [7].

### DR delays visual senescence by amplifying circadian rhythms in the eye

Metabolic dysfunction is strongly correlated with accelerated aging and eye-disease (e.g. diabetic retinopathy) [8, 9]. Declines in the circadian amplitude of clocks within the eye have been reported in wild-type mice with age and in models of diabetic retinopathy, which may further exacerbate disease pathology [10, 11]. Calorie restriction protects against several age-related eye diseases—dry-eye disease, cataracts, and age-related macular degeneration [12]. Calorie restriction also has a neuroprotective effect in photoreceptors and retinal-ganglion cells with age [12]. To date, no studies have investigated whether calorie restriction enhances circadian amplitude within the eye or whether its benefits within the eye are dependent on the molecular clock. Our results in flies demonstrate that DR amplifies circadian rhythms within the eye and delays visual senescence in a CLK-dependent manner. Additionally, we identified the DR-sensitive CLK-output genes *Gβ76c*, *retinin*, and *sunglasses* and demonstrated that their knockdown in the eye accelerated visual declines, thus indicating that DR’s neuroprotective role in the eye functions mechanistically through the molecular clock.

Several age-associated morphological and physiological declines have been reported in circadian mutant mouse models [13]. The positive-limb of the core molecular clock in mice is comprised of the basic-helix-loop-helix transcription factors BMAL1 and CLOCK [14]. Mice harboring whole-body genetic knockouts of either BMAL1 or CLOCK develop cataracts and corneal inflammation with age [13]. Additionally, photoreceptor-specific (cone-cell, HRGP-Cre x *Bmal1* fl/fl) BMAL1 knockout mice display a significantly altered circadian transcriptome, a shift in the distribution of short vs medium wavelength opsins, and a reduction in photoreceptor cell viability with age [15, 16]. Consistently, our data demonstrates that diminishing CLK function in adult animals (post-development) is sufficient to drive eye aging in flies. However, there are important distinctions between the mechanism of phototransduction used by mammalian rod and cone photoreceptors, and what exists in the fly.

In mammals, light-activated rhodopsin in rod and cone photoreceptor neurons couples to, and inactivates, cyclic nucleotide gated channels, hyperpolarizing the cell [17]. This is distinctly different from what occurs in the fly, where light-activated rhodopsin couples to a TRP channel, which when activated depolarizes the cell [18]. However, in a third class of mammalian photoreceptors, the intrinsically-photosensitive retinal ganglion cells (ipRGCs), there is a nearly identical mechanism of phototransduction to *Drosophila* [19]. The ipRGCs play a role in non-image forming light sensation, effecting pupillary constriction and the entrainment of the central circadian clock to light. There is some evidence that eliminating *Bmal1* in mice (either specifically in their ipRGCs or throughout their entire body) impairs the functionality of the ipRGCs [20]. This is consistent with what we observed when we disrupted *clk* in the *Drosophila* photoreceptors. Together, this suggests that there may be a conserved mechanism through which circadian clocks mediate the health of photoreceptor cells.

An inability to adequately respond to light stress may underly the accelerated photoreceptor aging we observe when CLK function is diminished in the eye of adult flies. Chronic exposure to phototoxic wavelengths or strong ambient light intensities, as well as mutations in light adaptation proteins, elevates intracellular calcium ion concentrations that result in rapid photoreceptor degeneration [21, 22]. Pittendrigh’s “escape the light” hypothesis posits that circadian rhythms evolved as a means for cells/organisms to anticipate and manage the deleterious effects of daily light exposure [23, 24]. One of the key neuroprotective functions of intrinsic clocks within photoreceptors is their ability to modulate time-of-day sensitivities to light. Electroretinogram (ERG) recordings in both flies and mammals have revealed a circadian response pattern that peaks at night when luminescence is approximately one-billion-fold less than during the day, and this pattern in light sensitivity is abolished in in circadian mutants [25, 26]. Interestingly, exposing rats to a bout of intense light at night results in significantly greater photoreceptor damage and degeneration than when the same treatment is performed during the day, thus highlighting the physiologic importance of the clocks in suppressing light sensitivity during the day [27]. Our acrophase analyses revealed that circadian transcripts that promote photoreceptor activation (Ca^2+^ influx) reach peak expression during the dark-phase, while genes that terminate the phototransduction response (i.e., reducing rhodopsin mediated signaling) peak in anticipation of the light-phase (Fig. 1f and Supplementary Fig. 1j).

### The eye regulates longevity in Drosophila

With age, declines in tissue homeostasis and chronic activation of the immune system increases local and systemic inflammation, termed “inflammaging.” The deleterious effects from inflammaging exacerbate pre-existing aging phenotypes and reduce survival [28, 29]. Interestingly, partial inhibition of the primary immune response regulator, NFkappaB, extends lifespan in *Drosophila* [30]. Photoreceptor degeneration is a main source of inflammation within the mouse retina [31]. Here, our results demonstrate that diminishing neuronal CLK function and forcing photoreceptor degeneration significantly elevates systemic immune responses. Furthermore, we report dampened AMP expression in the bodies of Rh null lines, indicating that reductions in phototransduction coincide with reduced systemic inflammation in the fly. We have also found that flies reared on DR, which improves photoreceptor viability, displayed dampened immune responses in comparison to flies reared on AL. Interestingly, photoreceptor degeneration caused by light- or calcium-mediated excitotoxicity is primarily the result of necrotic cell death [21]. Forced photoreceptor necrosis also results in necrotic cell death of surrounding cells [32]. Given that cytosolic f-actin can drive the sterile immune response in the fly, it is possible that the increased systemic inflammation we report is due in part to elevations in necrosis [33]. However, future studies will be needed to elucidate how diet and circadian rhythms influence necrotic cell death in photoceptors, and the effect this has on the local niche.

Circadian disruption, achieved either genetically or via chronic circadian misalignment with the environment, is associated with reduced longevity [34, 35]. Long-lived humans (i.e., centenarians) display significantly improved behavioral rhythms compared to “normal” aging groups [36]. Inversely, studies in mice and flies have demonstrated that organisms that display arrhythmic, or non-24h rhythms, are significantly shorter-lived than those who display near 24h (wild-type) circadian rhythms [37, 38]. Furthermore, chronic phase-adjustments, as is common in shift workers, is associated with early aging phenotypes and reduced lifespan in both mice and flies [35, 39]. Interestingly, placing BMAL1 knockout mice on calorie restriction fails to extend their lifespan [40]; although, these mice lack BMAL1 expression in all tissues and throughout development. To our knowledge, our study is the first to demonstrate that disruptions to the photoreceptor, and in particular to the photoreceptor clocks, is sufficient to shorten lifespan.

### DR extends lifespan in part by maintaining photoreceptor homeostasis

A number of studies have previously investigated the effects of light exposure on lifespan in *Drosophila* [41]. These studies, however, have not simultaneously examined the influence of diet and the influence of the photoreceptor cells. Exposure to short-wavelength light (i.e., blue-light), which is especially phototoxic, reduces survival in worms and flies [22, 42]. Interestingly, housing flies in a 12:12 blue-light/dark cycle significantly shortens lifespan even when those flies lack photoreceptors [22]. This effect appears to be directly related to blue-light mediated neuronal cell death (i.e., extraocular blue light sensing). Our lab previously demonstrated that DR-mediated lifespan extension is completely abolished in flies reared in constant lighting conditions (LL), while flies reared on AL experienced only a minor decrease in lifespan in LL [43]. We previously attested that LL blocked DR’s lost ability to extend lifespan because it induces arrhythmicity. A new, alternative, hypothesis is that the LL-mediated lifespan shortening on DR was also a result of the phototoxic effects of LL, which force photoreceptors to rapidly degenerate. The observation that knocking down *ATPα* in the eye (a model of forced photoreceptor degeneration) also significantly reduced lifespan on DR, further supports the photoreceptor hypothesis. Furthermore, as discussed above, photoreceptors regulate the timing of their light-sensitivity through the molecular clock. Therefore, housing a fly in LL would likely render their photoreceptor clocks arrhythmic, and increase the photoreceptor cells’ susceptibility to phototoxic stress. Interestingly, chronic dim-light exposure at night also shortens lifespan in *Drosophila* [44]. Our results here, indicate that DR protects flies from the lifespan-shortening effects of photoreceptor activation.

Although, forced photoreceptor degeneration is sufficient to significantly reduce longevity on DR, we also demonstrate that DR protects against the lifespan-shortening effects of photoreceptor activation during a normal 12:12 LD cycle. Flies reared on AL, which display dampened circadian rhythms within the eye, were selectively sensitive to lethality from the optogenetic activation of the photoreceptors. Inversely, we report that white-eyed flies, which are highly susceptible to light-mediated retinal degeneration, only display lifespan extension from constant darkness when they are reared on AL. Consistent with this observation, Rh null flies (which have reduced photoreceptor activity) display proportionally larger increases in lifespan when maintained on AL vs. DR. Importantly, although Rh expression is enriched in the photoreceptor cells, it is also expressed in other populations of neurons. Therefore, we cannot be certain that the lifespan extensions we observe in Rh null flies is solely the result of diminished Rh levels in the photoreceptor cells.

---

**Supplemental Figure 1.**
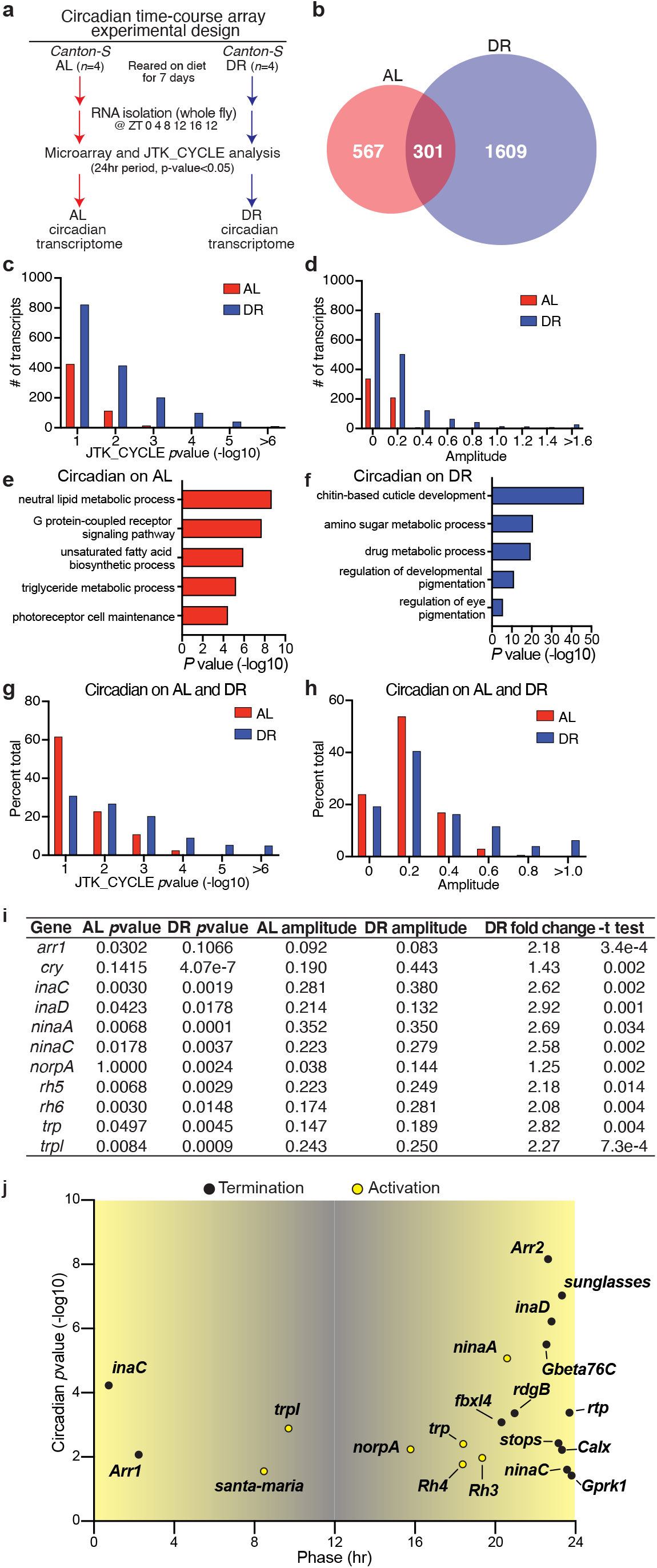
Dietary restriction amplifies circadian transcriptional output. (**a**) Experimental design of the time-course microarray and generation of the AL/DR circadian transcriptomes. Canton-S females were reared on AL or DR diets for 7 days. Flies were then collected, and mRNA was isolated from whole-fly lysates at 4-hour intervals for 24 hours (*n*=4 pooled mRNA samples from 30 flies per condition/timepoint). Circadian transcripts were identified with the JTK_CYCLE algorithm [45]. (**b**) Venn-diagram displaying the number of circadian transcripts that oscillate in flies reared on AL, DR, or in both diets. (**c-d**) Histograms of JTK_CYCLE *p*value statistics and circadian amplitudes of transcripts that cycle only on AL or DR diets. The y-axis indicates the total number of transcripts. A rightward shift was observed in *p*value and amplitude for transcripts that are circadian on DR compared to AL. (**e-f**) Gene-ontology enrichment categories corresponding to transcripts that cycle on AL (e) or DR (f). (**g-h**) Histograms of JTK_CYCLE *p*value statistics (g) and circadian amplitudes (h) of transcripts that are circadian on both diets. The y-axis indicates percent of transcripts out of the 301 total transcripts that oscillate on both diets. Transcripts that are circadian on both diets display smaller circadian *p*values and larger circadian amplitudes on DR compared to AL. (**i**) Table of phototransduction genes that are circadian on AL and DR. \**cry* is only circadian in DR, and *arr1* is only circadian on DR. Fold-changes and *t*test statistics were calculated by averaging the individual fold-changes in expression for each timepoint. (**j**) Circadian acrophase chart of transcripts that oscillate on DR and AL plotted as number of transcripts that peak at different timepoints throughout the day (as calculated by JTK_CYCLE algorithm). (**k**) Differential expression heatmap and associated GO terms for transcripts that are significantly upregulated (*n*=524) or downregulated (*n*=543) across all timepoints on DR compared to AL. Data were analyzed by student’s *t*test comparing expression values from AL and DR transcriptome from ZT0-20, and transcripts that display a *p*value less than 0.05 were considered differentially expressed.

**Supplemental Figure 2.**
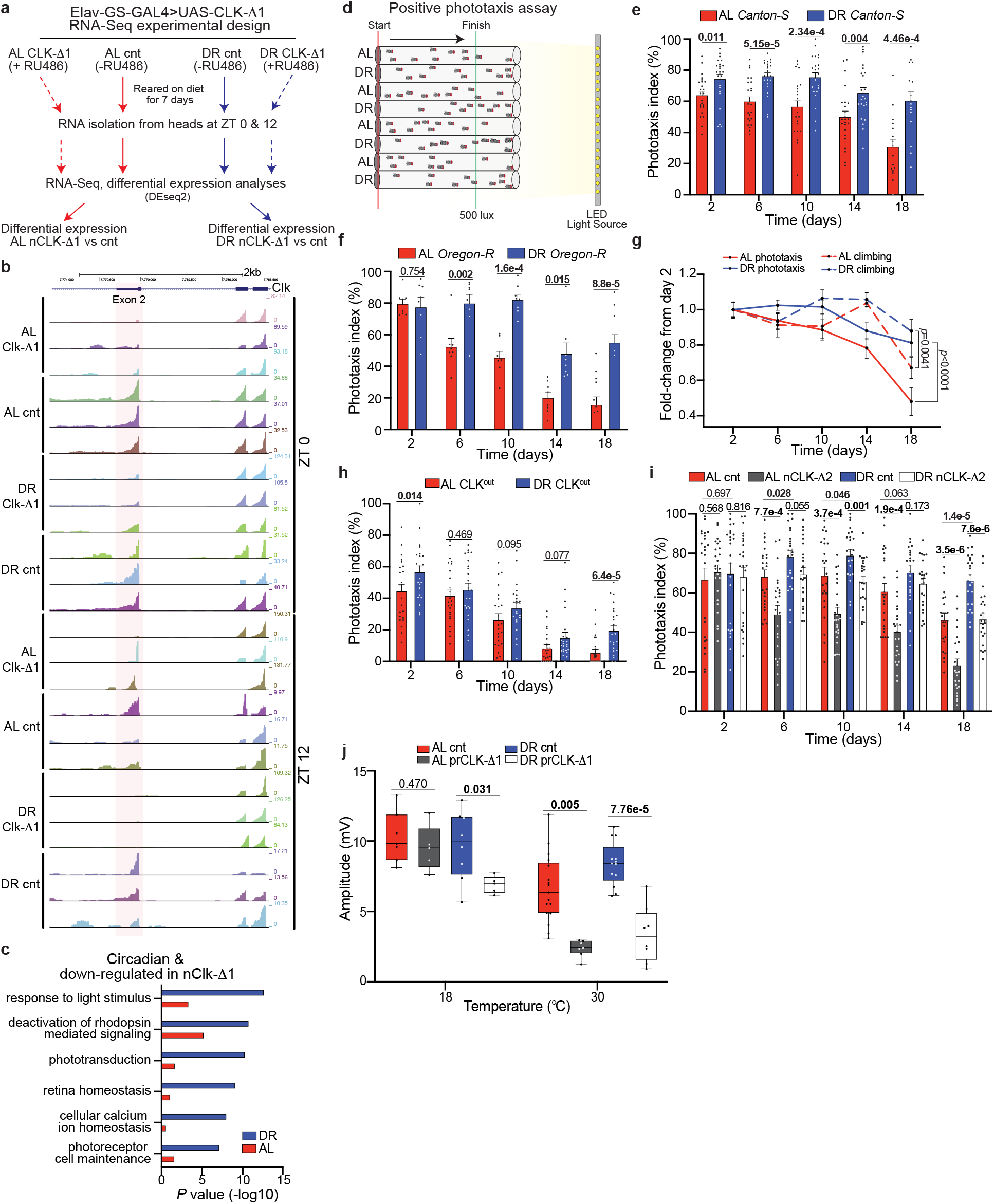
Design and additional analyses of nCLK-Δ1 RNA-Seq., positive phototaxis, and ERG experiments. (**a**) Design of nCLK-Δ1 RNA-Seq. Mated females were reared on AL or DR with the addition of vehicle or RU486 to induce the expression of CLK- Δ1 pan-neuronally for 7 days. mRNA was isolated from heads (*n*=3 biological replicates, *N*=30 heads per replicates) at ZT0 and ZT12. RNA-sequencing was performed and differentially expressed genes were identified with the DEseq2 software [46] package. (**b**) UCSC genome browser visualization of the individual tracks for each nCLK-Δ1 RNA-Seq sample zoomed into exon 2 of *clk* (chr3L:7,766,807-7,773,169). Exon 2 (highlighted in red) of *clk* encodes the basic helix-loop-helix domain (DNA binding) of CLK that is selectively ablated in CLK-Δ1 flies. Overexpression of CLK-Δ1 results in a relative decrease in the ratio of tags at exon 2 vs exon 3-4 (right), while exon 3-4 display elevated tag density compared to control samples. Track size is normalized for each sample and the total number of tags is indicated as the top number (color coded to match each track) on the far right. (**c**) Geneontology enrichment terms and *p*value statistics for genes that are circadian in young heads (GEO81100) [7] and significantly down-regulated in the nCLKΔ1 RNA-Seq on AL or DR. (**d**) Diagram of positive-phototaxis setup. Flies are sorted in clear elongated fly vials, dark adapted for 15 minutes, knocked to the bottom of the vial, and then laid horizontally and perpendicular to an LED light source. Once the light is turned on flies that reach the green line are scored as “positive-phototaxis” and counted at 15, 30, 45 seconds (See methods for additional details). (**e**) Positive phototaxis responses for *Canton-S* females reared on AL or DR diets. See methods for calculation of phototaxis index. For each timepoint results are represented as average percent positive phototaxis +/- SEM (*n*=24 biological reps, *N*=480 flies per condition). (**f**) Phototaxis responses for *Oregon-R* females. For each timepoint results are represented as average percent phototaxis response +/- SEM (*n*=8 biological replicates, *N*=160 flies per condition). (**g**) *Canton-S* climbing activity and positive phototaxis plotted as fold-change from responses at day 2. (*n*=24 biological replicates, *N*=480 flies per condition). (**h**) Positive phototaxis responses for *Clk^ou^*^t^ females reared on AL or DR diets. For each timepoint results are represented as average percent positive phototaxis +/- SEM (*n*=24 biological reps, *N*=480 flies per condition). (**i**) Positive phototaxis responses for nCLK-Δ2 flies (Elav-GS-GAL4>UAS-CLK-Δ2. For each timepoint results are represented as average percent positive phototaxis +/- SEM (*n*=24 biological replicates, *N*=480 flies per condition). (**j**) Box-plots of electroretinogram amplitudes for prCLK-Δ1 (Trpl-GAL4;GAL80>UAS-CLK-Δ1^OC^) and control flies (Trpl- GAL4;GAL80>*CantonS*^OC^) reared at 18°C (GAL80 active, GAL4 inactive) and 30°C (GAL80 inactive, GAL4 active) for 6 days. Illuminance was set at 15000 Lux. (e-f and h-j) *P*values were determined by two-tailed Student’s *t*test (unpaired) at each timepoint. (g) *P*values were determined by two-tailed Student’s *t*test (unpaired) across genotypes.

**Supplemental Figure 3.**
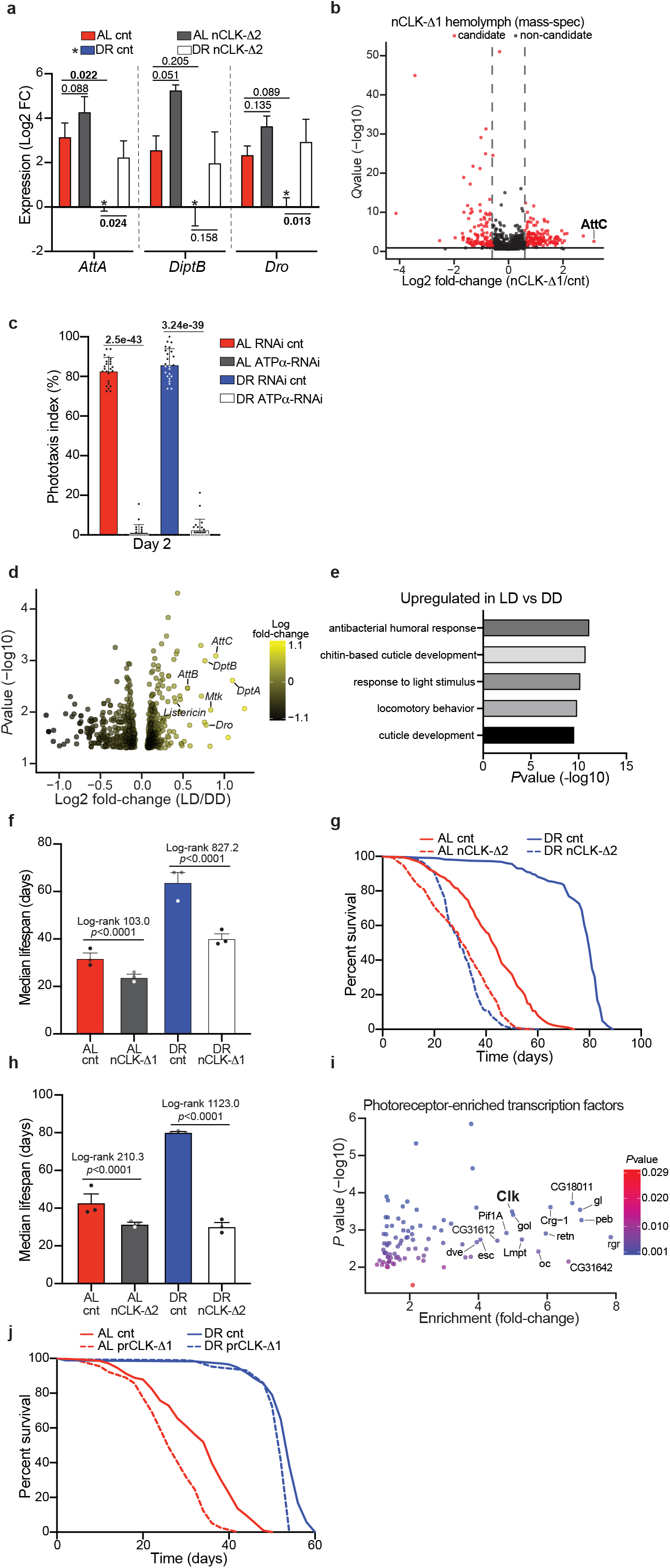
nCLK elevates immune responses and shortens longevity in a diet-dependent fashion. (**a**) Relative expression of AMP genes (*AttA*, *DiptB*, and *Dro*) calculated by RT-qPCR with mRNA isolated from nCLK-Δ2 bodies. Results are plotted as average Log2 fold-change in expression calculated by the ΔΔ-Ct method, normalized to DR vehicle treated control samples as well as *rp49* +/- SEM (*n*=3 biological replicates, *N*=30 flies per biological replicate). (**b**) Volcano-plot of hemolymph proteins identified by tandem mass-spectrometry comparing nCLK-Δ1 (RU486 treated, *N*=300) and control (vehicle treated, *N*=300) flies reared on AL at day 14. Each dot represents an individual protein with a statistical significance less than 0.0001 comparing nCLK-Δ1 and control hemolymph samples. Black dots are considered to be differentially expressed protein candidates with a Log2 fold-change cutoff of ± 0.6. AttC was the most highly up-regulated protein in nCLK-Δ1 hemolymph compared to control. (**c**) Volcano-plot of gene expression changes in heads of *y.w.* flies housed in 12:12 LD vs constant darkness (DD) from Wijnen et al, 2006 (GSE3842) [47]. Fold-changes in response to light were calculated by averaging the changes in expression at each timepoint from a circadian time-course microarray (ZT 2, 6, 10, 14, 18, 22) and comparing expression between flies housed in LD compared to DD. (**d**) The top-5 enriched gene-ontology categories corresponding to genes that are upregulated in heads of flies housed in LD vs DD. (**e**) Positive phototaxis responses with eye-specific knockdown of *ATPα* (GMR-GAL4>UAS-*ATPα*-RNAi) compared to RNAi control flies (GMR-GAL4>UAS-*mCherry*-RNAi). For each timepoint results are represented as average phototaxis response +/- SEM (*n*=24 biological replicates, *N*=480 flies per condition). (**f**) Median lifespan of nCLK-Δ1 flies corresponding to lifespans in (Fig. 3D). Data are plotted as the average median lifespan of the 3 biological replicates and error bars indicate +/- SEM. (**g**) Survival analysis of nCLK-Δ2 flies. Survival data is plotted as an average of three independent lifespan repeats. Control flies (vehicle treated): AL *N*=505, DR *N*=504; nCLK-Δ1 flies (RU486 treated): AL *N*=497, DR *N*=508. (**h**) Median lifespan of nCLK-Δ2 flies corresponding to lifespans in (Supplemental Fig. 3D). Data are plotted as the average median lifespan of the 3 biological replicates and error bars indicate +/- SEM. (**i**) Volcano-plot of photoreceptor-specific transcription factors [48]. (**j**) Survival analysis of prCLK-Δ1 flies. Survival data is plotted from one independent lifespan. Control flies (Trpl-GAL4;GAL80>UAS-*CantonS*^OC^): AL *N*=206, DR *N*=172; prCLK-Δ1 flies (Trpl-GAL4;GAL80> UAS-CLK-Δ1^OC^): AL *N*=154, DR *N*=190. *The *CantonS* and UAS-CLK-Δ1 parental lines were outcrossed to w^1118^ and the F1 generations share the same genetic background. (a) *P*values were calculated with the pairwise Student’s *t*test comparing Log2 fold-changes in expression. (e) *P*values were determined by two-tailed Student’s *t*test (unpaired) at each timepoint. (f, h) *P*values were determined by Chi square from Log-rank (Mantel-Cox) test.

**Supplemental Figure 4.**
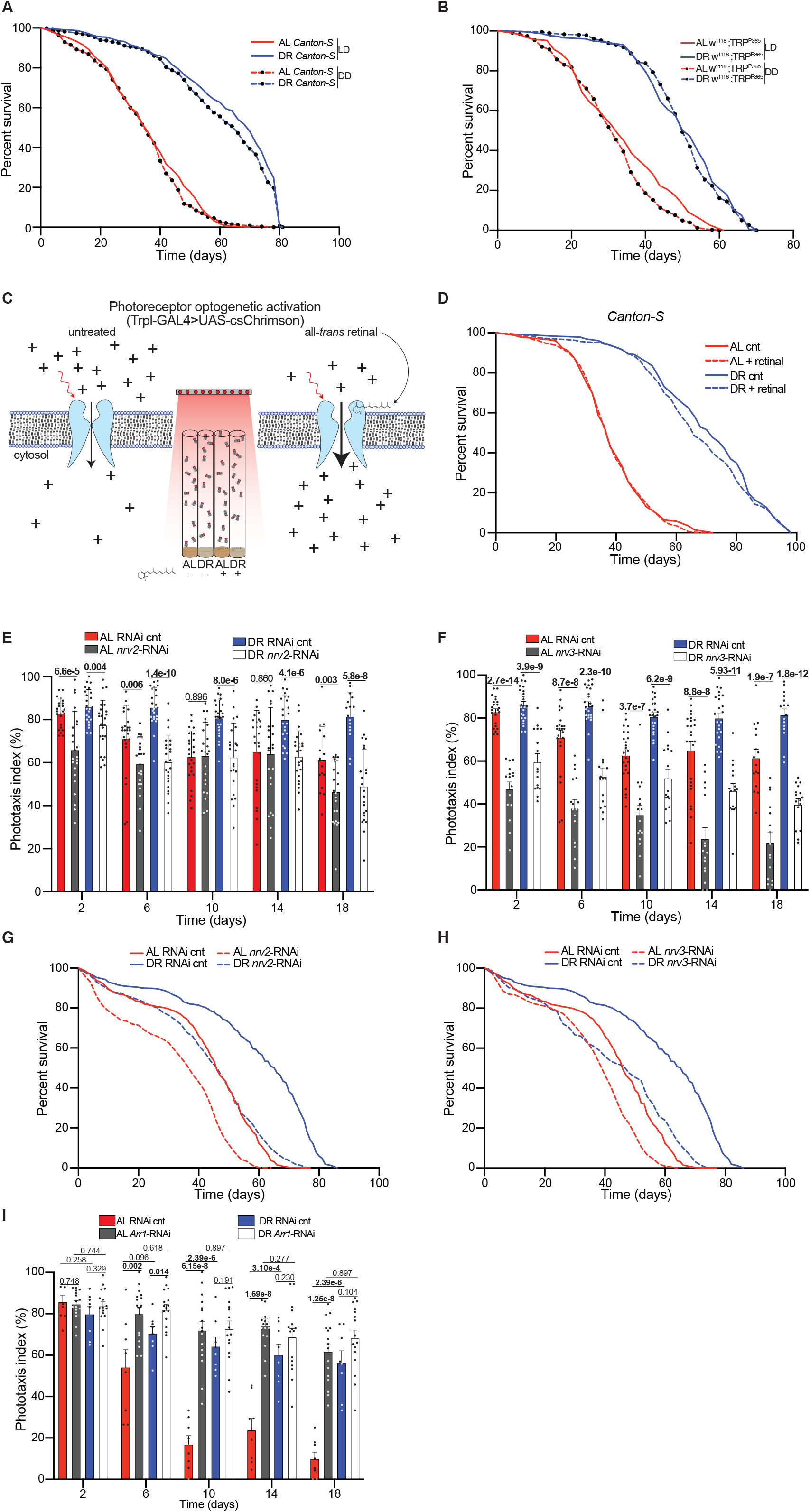
Lighting and retinal control lifespans, optogenetic activation diagram, and lifespans of RNAi-mediated knockdown of *ATPα*-subunits in the eye. (**a**) Survival analysis of *Canton-S* wildtype flies housed in 12:12h LD and constant darkness (DD). Survival data is plotted as an average of three independent lifespan crosses. AL LD *N*=549, AL DD *N*=510, DR LD *N*=558, DR DD *N*=509. (**b**) Survival analysis of white-eyed, photoreceptor null flies (*w^1118^*; TRP^P365^) housed in 12:12h LD or DD. Survival data is plotted as an average of two independent lifespan repeats. LD housed flies: AL *N*=290, DR *N*=373; DD housed flies: AL *N*=301, DR *N*=357. (**c**) Diagram of optogentic activation of photoreceptors. The photoreceptor-specific driver, Trpl-GAL4, drives the expression of a red-shifted csChrimson channel in R1-R8 photoreceptors. Addition of all-*trans* retinal (50μM) in the fly media promotes the opening of optogenetic channels in the presence of red-light, allowing the flow of positively charged ions into the cytosol to activate photoreceptors. (**d**) Survival analysis of *Canton-S* flies reared in 12:12 red-light:dark on AL and DR diets with the addition of all-*trans* retinal or vehicle (control). Survival data is plotted as an average of two independent lifespan repeats. All-*trans* retinal treated flies: AL *N*= 340, DR *N*=328; Vehicle treated flies: AL *N*=347, DR *N*=328. (**e-f**) Positive phototaxis responses with eye-specific knockdown *nrv2* (e, GMR-GAL4>UAS-*nrv2*-RNAi), and *nrv3* (f, GMR-GAL4>UAS-*nrv3*-RNAi) compared to RNAi control flies (GMR-GAL4>UAS-*mCherry*-RNAi). For each timepoint results are represented as average phototaxis response +/- SEM (RNAi control and *nrv2*: *n*=24 biological replicates, *N*=480 flies per condition; *nrv3*: *n*=16 biological replicates, *N*=320 flies per condition). (**g-h**) Survival analysis of eye-specific *nrv2* (g) and *nrv3* (h) RNAi knockdown flies compared to RNAi control flies. Survival data is plotted as an average of three independent lifespan repeats for RNAi controls and *nrv2* RNAi flies, and two independent lifespan crosses for *nrv3* RNAi knockdown flies. RNAi cnt flies: AL *N*=493, DR *N*=490; *nrv2* RNAi flies: AL *N*=482, DR *N*=513; *nrv3* RNAi flies: AL *N*=301, DR *N*=288. (e-f) *P*values were determined by two-tailed Student’s *t*test (unpaired) at each timepoint.

**Supplemental Figure 5.**
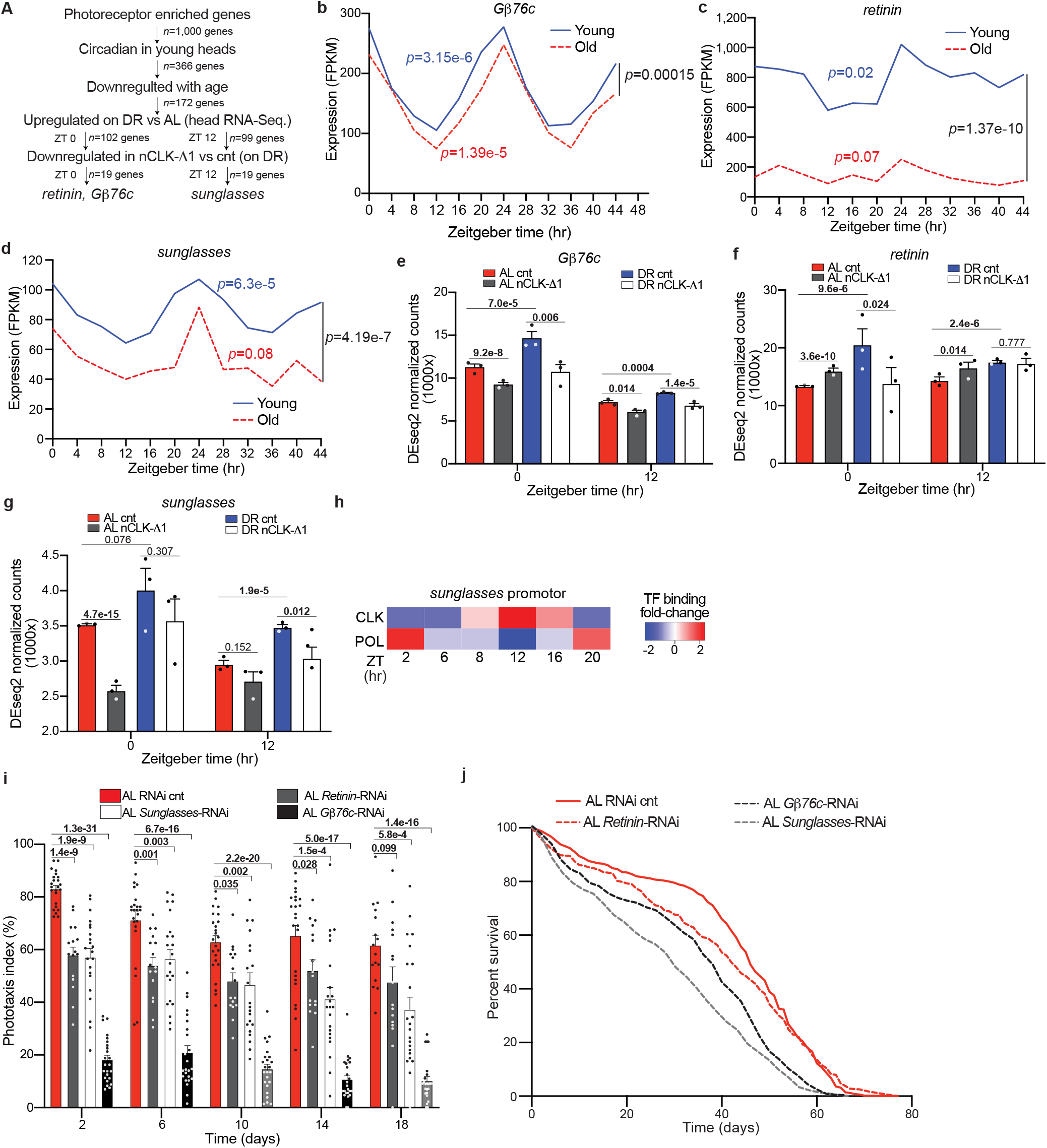
Identification of photoreceptor enriched CLK-output genes, and additional analyses with eye-specific knockdown of *Gβ76c*, *retinin*, and *sunglasses*. (**a**) Bioinformatics pipeline for identifying photoreceptor enriched, CLK-output genes. (**b-d**) Circadian expression of *Gβ76c*, *retinin*, and *sunglasses* and their corresponding circadian *p*value statistics for young (5-day old) and old (55-day old) wildtype heads from Kuintzle *et al*., 2017 [7]. (**e-g**) Normalized expression counts for *Gβ76c*, *retinin*, and *sunglasses* from the nCLK-Δ1 RNA-Seq. Results are represented as average expression counts calculated by DEseq2 +/- SEM. (**h**) Heatmap of CLK and POL (*Drosophila* polymerase) tag-densities at the 5’-untranslated region of the *sunglasses* promoter over a circadian time-course from ChIP-Chip analyses [49]. Consistent with other direct CLK target genes, Abruzzi *et al*., 2011 report maximal CLK binding at ZT 12, while POL displayed antiphasic binding to that of CLK and aligned with the phase of *sunglasses* mRNA expression (ZT 0-2). *CLK binding was not observed in GMR-HID heads suggesting *sunglasses* is under CLK transcriptional regulation specifically in the neurons of the eye. (**i**) Positive phototaxis responses with eye-specific knockdown of *Gβ76c* (GMR-GAL4>UAS-*Gβ76c*-RNAi), *retinin* (GMR-GAL4>UAS-*retinin*-RNAi), and *sunglasses* (GMR-GAL4>UAS-*sunglasses*-RNAi) compared to RNAi control flies (GMR-GAL4>UAS-*mCherry*-RNAi) reared on AL. For each timepoint results are represented as average phototaxis response +/- SEM (RNAi control *n*=24 biological replicates, *N*=480 flies per condition; *Gβ76c* RNAi *n*=24 biological replicates, *N*=480 flies per condition, *retinin* RNAi *n*=16 biological replicates, *N*=384 flies per condition; *sunglasses* RNAi *n*=24 biological replicates, *N*=480 flies per condition). (**j**) Survival analysis of eye-specific *Gβ76c*-RNAi, *retinin*-RNAi, *sunglasses*-RNAi, and RNAi control knockdown flies compared to RNAi control flies reared on AL. Survival data is plotted as an average of three independent lifespan repeats for RNAi control, *Gβ76c*-RNAi, sunglasses-RNAi flies and two independent lifespan repeats for *retinin-*RNAi flies. RNAi cnt flies: *N*=493; *Gβ76c* RNAi flies: *N*=543; *retinin* RNAi flies: *N*=353; *sunglasses* RNAi flies: *N*=503. (b-d) Circadian *p*values were determined by ARSER algorithm by Kuintzle *et al*., 2017 [7] (AL=red, DR=blue). To compare gene expression profiles with age we utilized the two-tailed Student’s *ttest* (paired) to determine *P*values (black). (e-g) *P*values were determined by DEseq2 differential expression analysis. (i) *P*values were determined by two-tailed Student’s *t*test (unpaired) at each timepoint comparing the phototaxis index of RNAi control flies to *retinin*- and *sunglasses*-RNAi flies.

**Supplemental Table 1.**
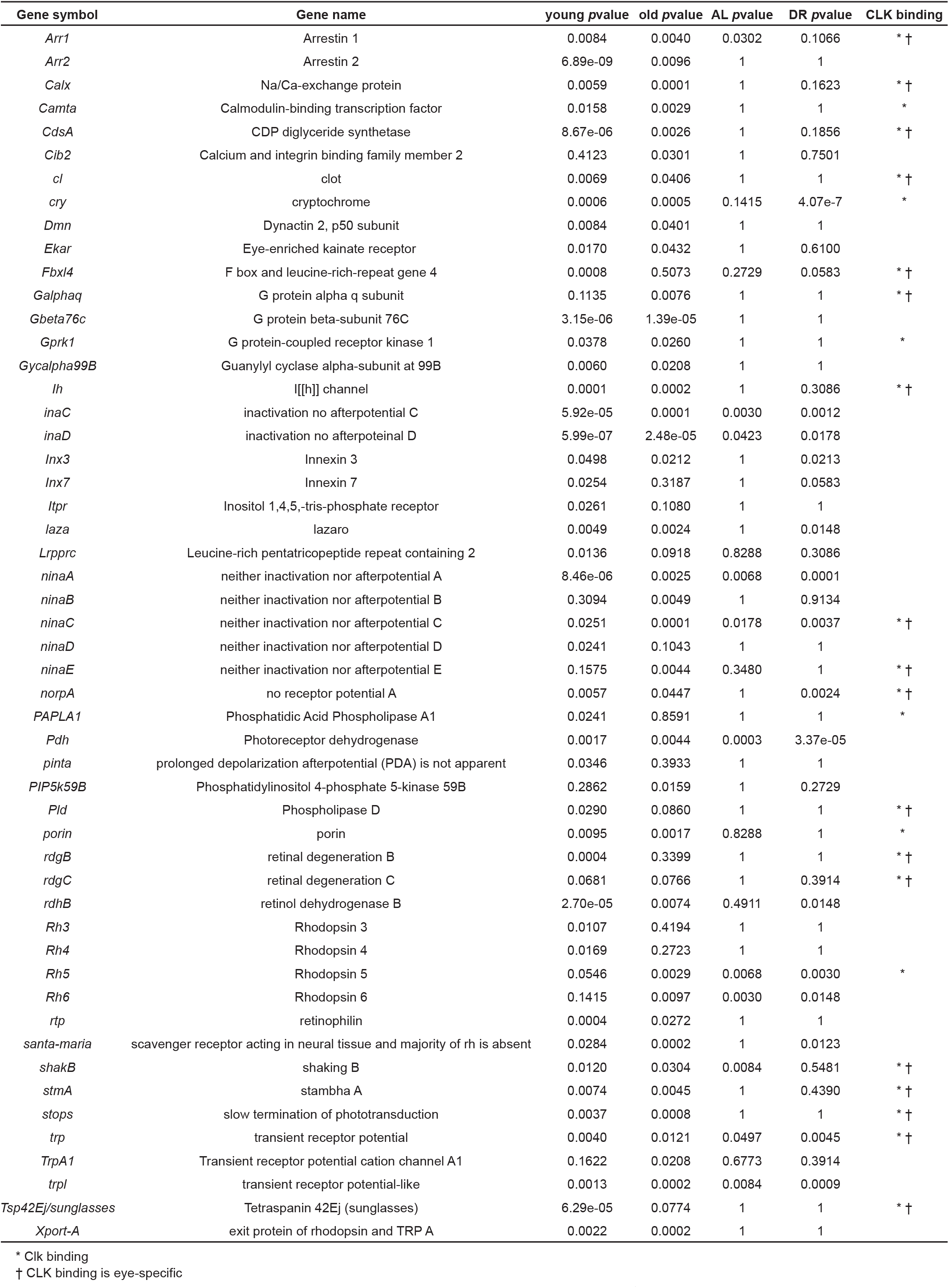
Circadian statistics and CLK binding of light-response genes. Circadian pvalue statistics for light response genes in young and old wildtype heads (calculated by ARSER by Kuintzle et al., 2017 [7]) and whole flies reared on AL and DR (calculated by JTK_CYCLE in this study).

**Supplemental Table 2.**
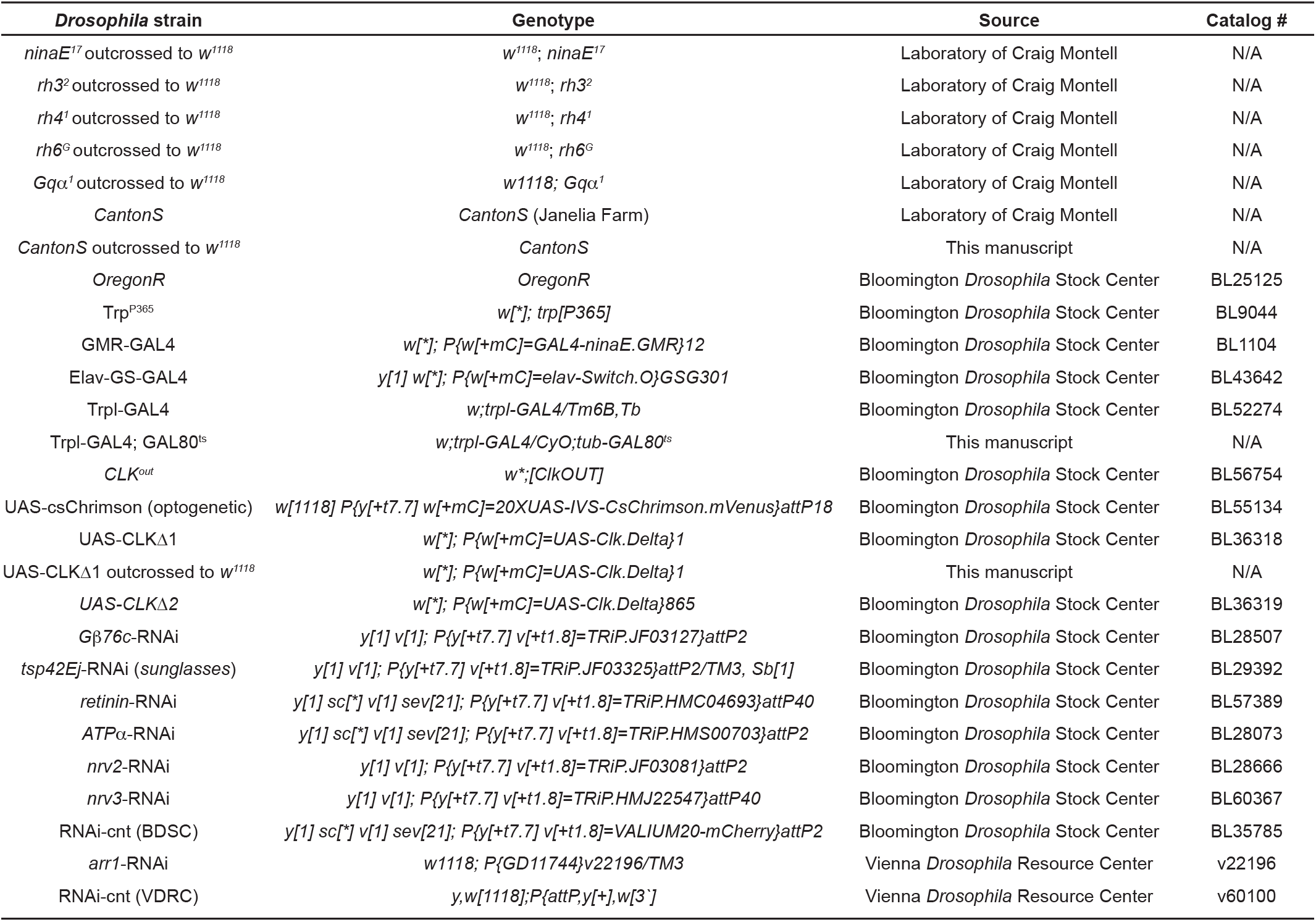
*Drosophila* strains used in this study.

**Supplemental Data 1. AL and DR circadian transcriptome analyses**. These files contain JTK_CYCLE statistics accompanied gene-ontology enrichment terms/scores for AL and DR circadian transcriptomes, circadian acrophase analyses, and differential gene-expression analyses.

**Supplemental Data 2. Gene-ontology enrichment analyses of genes that are circadian in young heads.** These files contain the enriched biological processes in young wild-type heads from Kuintzle et al., 2017, highlighting circadian processes within the eye.

**Supplemental Data 3. Additional nCLK-Δ1 RNA-Seq analyses.** Included in these files are normalized count reads generated by DEseq2 across all experimental groups and replicates from the nCLK-Δ1 RNA-Seq. The normalized expression counts across all samples for the gene-ontology terms “Deactivation of rhodopsin mediated signaling” and “Antimicrobial humoral response” (corresponding to Fig. 2b and 3b) are also reported.

**Supplemental Data 4. Cross-comparison of wild-type circadian transcriptome and nCLK-Δ1 RNA-Seq analyses.** These files include gene-ontology enrichment terms/scores for genes that are circadian in wild-type heads (from Kuintzle *et al.*, 2017, GSE81100) and downregulated in nCLK-Δ1 heads.

**Supplemental Data 5. nCLK-Δ1 hemolymph mass-spec analysis.** These files contain the proteins identified and quantification of differential expression comparing proteomic profiles between nCLK-Δ1 and control hemolymph. Enriched bioprocesses are also included for significantly up- or downregulated proteins.

**Supplemental Data 6. Bioinformatic pipeline for identification of eye-specific and DR- sensitive CLK-output genes *Gbeta76c*, *retinin*, and *sunglasses*.** These files provide the filtered gene-lists that correspond to the bioinformatic filtering steps performed in Fig. 5a-d and Supplementary Fig. 5a.

**Supplemental Data 7. Survival analyses.** These files report lifespan statistics (Log-Rank and Hazard Ratios) and group sizes (*n*) for the survival analyses performed.

**Supplemental Data 8. Analyses of transcriptional responses to light.** These files report gene-ontology enrichment terms/scores for genes that are differentially expressed in wild-type fly heads in response to being housed in 12:12h LD vs constant dark (from Wijnen *et al*., 2006, GSE3842) and correspond to Supplementary Fig. 3D-E.

**Supplemental Data 9. Positive phototaxis responses and statistics.** These files report detailed statistics (*t*test and 2way ANOVA) for the positive phototaxis experiments performed.

**Supplemental Data 10. Electroretinogram analyses and statistics**. These files include detailed *t*test statistics for the ERG assays performed at day 14 and 21 in nCLK-Δ1 flies.

**Supplemental Data 11. Experimental materials.**

## References

1. Rijo-Ferreira, F. and J.S. Takahashi, Genomics of circadian rhythms in health and disease. Genome Med, 2019. 11(1): p. 82.

2. Patke, A., M.W. Young, and S. Axelrod, Molecular mechanisms and physiological importance of circadian rhythms. Nat Rev Mol Cell Biol, 2020. 21(2): p. 67–84.

3. Rouyer, F., Clock genes: from Drosophila to humans. Bull Acad Natl Med, 2015. 199(7): p. 1115–1131.

4. Ogueta, M., R.C. Hardie, and R. Stanewsky, Non-canonical Phototransduction Mediates Synchronization of the Drosophila melanogaster Circadian Clock and Retinal Light Responses. Curr Biol, 2018. 28(11): p. 1725–1735 e3.

5. Hall, H., et al., Transcriptome profiling of aging Drosophila photoreceptors reveals gene expression trends that correlate with visual senescence. BMC Genomics, 2017. 18(1): p. 894.

6. Lin, J.B., K. Tsubota, and R.S. Apte, A glimpse at the aging eye. NPJ Aging Mech Dis, 2016. 2: p. 16003.

7. Felder-Schmittbuhl, M.P., et al., Ocular Clocks: Adapting Mechanisms for Eye Functions and Health. Invest Ophthalmol Vis Sci, 2018. 59(12): p. 4856–4870.

8. Baba, K. and G. Tosini, Aging Alters Circadian Rhythms in the Mouse Eye. J Biol Rhythms, 2018. 33(4): p. 441–445.

9. Kapahi, P., M. Kaeberlein, and M. Hansen, Dietary restriction and lifespan: Lessons from invertebrate models. Ageing Res Rev, 2017. 39: p. 3–14.

10. Katewa, S.D., et al., Peripheral Circadian Clocks Mediate Dietary Restriction-Dependent Changes in Lifespan and Fat Metabolism in Drosophila. Cell Metab, 2016. 23(1): p. 143–54.

11. Eckel-Mahan, K.L., et al., Reprogramming of the circadian clock by nutritional challenge. Cell, 2013. 155(7): p. 1464–78.

12. Honma, T., et al., High-fat diet intake accelerates aging, increases expression of Hsd11b1, and promotes lipid accumulation in liver of SAMP10 mouse. Biogerontology, 2012. 13(2): p. 93–103.

13. Montell, C., Visual transduction in Drosophila. Annu Rev Cell Dev Biol, 1999. 15: p. 231–68.

14. Montell, C., Drosophila visual transduction. Trends Neurosci, 2012. 35(6): p. 356–63.

15. Kuintzle, R.C., et al., Circadian deep sequencing reveals stress-response genes that adopt robust rhythmic expression during aging. Nat Commun, 2017. 8: p. 14529.

16. Spitschan, M., et al., Variation of outdoor illumination as a function of solar elevation and light pollution. Sci Rep, 2016. 6: p. 26756.

17. Gu, Y., et al., Mechanisms of light adaptation in Drosophila photoreceptors. Curr Biol, 2005. 15(13): p. 1228–34.

18. Oberwinkler, J. and D.G. Stavenga, Calcium transients in the rhabdomeres of dark- and light-adapted fly photoreceptor cells. J Neurosci, 2000. 20(5): p. 1701–9.

19. Shieh, B.H., Molecular genetics of retinal degeneration: A Drosophila perspective. Fly (Austin), 2011. 5(4): p. 356–68.

20. Voolstra, O. and A. Huber, Ca(2+) Signaling in Drosophila Photoreceptor Cells. Adv Exp Med Biol, 2020. 1131: p. 857–879.

21. Katewa, S.D. and P. Kapahi, Dietary restriction and aging, 2009. Aging Cell, 2010. 9(2): p. 105–12.

22. Hotta, Y. and S. Benzer, Abnormal electroretinograms in visual mutants of Drosophila. Nature, 1969. 222(5191): p. 354–6.

23. Vilinsky, I. and K.G. Johnson, Electroretinograms in Drosophila: a robust and genetically accessible electrophysiological system for the undergraduate laboratory. J Undergrad Neurosci Educ, 2012. 11(1): p. A149–57.

24. Garschall, K. and T. Flatt, The interplay between immunity and aging in Drosophila. F1000Res, 2018. 7: p. 160.

25. Pletcher, S.D., et al., Genome-wide transcript profiles in aging and calorically restricted Drosophila melanogaster. Curr Biol, 2002. 12(9): p. 712–23.

26. Ferrandon, D., et al., The Drosophila systemic immune response: sensing and signalling during bacterial and fungal infections. Nat Rev Immunol, 2007. 7(11): p. 862–74.

27. Handke, B., et al., The hemolymph proteome of fed and starved Drosophila larvae. PLoS One, 2013. 8(6): p. e67208.

28. Palladino, M.J., et al., Neural dysfunction and neurodegeneration in Drosophila Na+/K+ ATPase alpha subunit mutants. J Neurosci, 2003. 23(4): p. 1276–86.

29. Baumann, O., *Distribution of Na+,*K(+)-ATPase in photoreceptor cells of insects. Int Rev Cytol, 1997. 176: p. 307–48.

30. Damulewicz, M., E. Rosato, and E. Pyza, Circadian regulation of the Na+/K+-ATPase alpha subunit in the visual system is mediated by the pacemaker and by retina photoreceptors in Drosophila melanogaster. PLoS One, 2013. 8(9): p. e73690.

31. Luan, Z., K. Reddig, and H.S. Li, Loss of Na(+)/K(+)-ATPase in Drosophila photoreceptors leads to blindness and age-dependent neurodegeneration. Exp Neurol, 2014. 261: p. 791–801.

32. Wijnen, H., et al., Control of daily transcript oscillations in Drosophila by light and the circadian clock. PLoS Genet, 2006. 2(3): p. e39.

33. Kumar, J.P., Building an ommatidium one cell at a time. Dev Dyn, 2012. 241(1): p. 136–49.

34. Wang, T. and C. Montell, Phototransduction and retinal degeneration in Drosophila. Pflugers Arch, 2007. 454(5): p. 821–47.

35. Kurada, P. and J.E. O’Tousa, Retinal degeneration caused by dominant rhodopsin mutations in Drosophila. Neuron, 1995. 14(3): p. 571–9.

36. Li, Q., et al., Temperature and Sweet Taste Integration in Drosophila. Curr Biol, 2020. 30(11): p. 2051–2067 e5.

37. Vasiliauskas, D., et al., Feedback from rhodopsin controls rhodopsin exclusion in Drosophila photoreceptors. Nature, 2011. 479(7371): p. 108–12.

38. Leung, N.Y., et al., Functions of Opsins in Drosophila Taste. Curr Biol, 2020. 30(8): p. 1367–1379 e6.

39. Shostal, O.A. and A.A. Moskalev, The genetic mechanisms of the influence of the light regime on the lifespan of Drosophila melanogaster. Front Genet, 2012. 3: p. 325.

40. Nash, T.R., et al., Daily blue-light exposure shortens lifespan and causes brain neurodegeneration in Drosophila. NPJ Aging Mech Dis, 2019. 5: p. 8.

41. Ferreiro, M.J., et al., Drosophila melanogaster White Mutant w(1118) Undergo Retinal Degeneration. Front Neurosci, 2017. 11: p. 732.

42. Yoon, J., et al., Novel mechanism of massive photoreceptor degeneration caused by mutations in the trp gene of Drosophila. J Neurosci, 2000. 20(2): p. 649–59.

43. Scott, K., et al., Gq alpha protein function in vivo: genetic dissection of its role in photoreceptor cell physiology. Neuron, 1995. 15(4): p. 919–27.

44. Shieh, B.H., I. Kristaponyte, and Y. Hong, Distinct roles of arrestin 1 protein in photoreceptors during Drosophila development. J Biol Chem, 2014. 289(26): p. 18526–34.

45. Satoh, A.K. and D.F. Ready, Arrestin1 mediates light-dependent rhodopsin endocytosis and cell survival. Curr Biol, 2005. 15(19): p. 1722–33.

46. Klapoetke, N.C., et al., Independent optical excitation of distinct neural populations. Nat Methods, 2014. 11(3): p. 338–46.

47. Simpson, J.H. and L.L. Looger, Functional Imaging and Optogenetics in Drosophila. Genetics, 2018. 208(4): p. 1291–1309.

48. Dolph, P.J., et al., An eye-specific G beta subunit essential for termination of the phototransduction cascade. Nature, 1994. 370(6484): p. 59–61.

49. Stahl, A.L., et al., The cuticular nature of corneal lenses in Drosophila melanogaster. Dev Genes Evol, 2017. 227(4): p. 271–278.

50. Kryuchkov, M., et al., Reverse and forward engineering of Drosophila corneal nanocoatings. Nature, 2020. 585(7825): p. 383–389.

51. Xu, H., et al., A lysosomal tetraspanin associated with retinal degeneration identified via a genome-wide screen. EMBO J, 2004. 23(4): p. 811–22.

52. Abruzzi, K.C., et al., Drosophila CLOCK target gene characterization: implications for circadian tissue-specific gene expression. Genes Dev, 2011. 25(22): p. 2374–86.

53. Patel, S.A., et al., Circadian clocks govern calorie restriction-mediated life span extension through BMAL1- and IGF-1-dependent mechanisms. FASEB J, 2016. 30(4): p. 1634–42.

54. Giebultowicz, J.M. and D.M. Long, Ageing and Circadian rhythms. Curr Opin Insect Sci, 2015. 7: p. 82–86.

55. Hood, S. and S. Amir, The aging clock: circadian rhythms and later life. J Clin Invest, 2017. 127(2): p. 437–446.

56. Boomgarden, A.C., et al., Chronic circadian misalignment results in reduced longevity and large-scale changes in gene expression in Drosophila. BMC Genomics, 2019. 20(1): p. 14.

57. Martinez-Nicolas, A., et al., Circadian monitoring as an aging predictor. Sci Rep, 2018. 8(1): p. 15027.

58. Retaux, S. and D. Casane, Evolution of eye development in the darkness of caves: adaptation, drift, or both? Evodevo, 2013. 4(1): p. 26.

59. Lang, S., et al., A conserved role of the insulin-like signaling pathway in diet-dependent uric acid pathologies in Drosophila melanogaster. PLoS Genet, 2019. 15(8): p. e1008318.

60. Nicholson, L., et al., Spatial and temporal control of gene expression in Drosophila using the inducible GeneSwitch GAL4 system. I. Screen for larval nervous system drivers. Genetics, 2008. 178(1): p. 215–34.

61. Katewa, S.D., et al., Intramyocellular fatty-acid metabolism plays a critical role in mediating responses to dietary restriction in Drosophila melanogaster. Cell Metab, 2012. 16(1): p. 97–103.

62. Bolger, A.M., M. Lohse, and B. Usadel, Trimmomatic: a flexible trimmer for Illumina sequence data. Bioinformatics, 2014. 30(15): p. 2114–20.

63. Kim, D., B. Langmead, and S.L. Salzberg, HISAT: a fast spliced aligner with low memory requirements. Nat Methods, 2015. 12(4): p. 357–60.

64. Liao, Y., G.K. Smyth, and W. Shi, featureCounts: an efficient general purpose program for assigning sequence reads to genomic features. Bioinformatics, 2014. 30(7): p. 923–30.

65. Love, M.I., W. Huber, and S. Anders, Moderated estimation of fold change and dispersion for RNA-seq data with DESeq2. Genome Biol, 2014. 15(12): p. 550.

66. Wang, S., et al., The retromer complex is required for rhodopsin recycling and its loss leads to photoreceptor degeneration. PLoS Biol, 2014. 12(4): p. e1001847.

67. Wes, P.D., et al., Termination of phototransduction requires binding of the NINAC myosin III and the PDZ protein INAD. Nat Neurosci, 1999. 2(5): p. 447–53.

68. Vang, L.L., A.V. Medvedev, and J. Adler, Simple ways to measure behavioral responses of Drosophila to stimuli and use of these methods to characterize a novel mutant. PLoS One, 2012. 7(5): p. e37495.

69. Christensen, D.G., et al., Identification of Novel Protein Lysine Acetyltransferases in Escherichia coli. mBio, 2018. 9(5).

70. Collins, B.C., et al., Multi-laboratory assessment of reproducibility, qualitative and quantitative performance of SWATH-mass spectrometry. Nat Commun, 2017. 8(1): p. 291.

71. Schilling, B., B.W. Gibson, and C.L. Hunter, Generation of High-Quality SWATH((R)) Acquisition Data for Label-free Quantitative Proteomics Studies Using TripleTOF((R)) Mass Spectrometers. Methods Mol Biol, 2017. 1550: p. 223–233.

72. Charlton-Perkins, M.A., et al., Multifunctional glial support by Semper cells in the Drosophila retina. PLoS Genet, 2017. 13(5): p. e1006782.

73. Hughes, M.E., J.B. Hogenesch, and K. Kornacker, JTK_CYCLE: an efficient nonparametric algorithm for detecting rhythmic components in genome-scale data sets. J Biol Rhythms, 2010. 25(5): p. 372–80.

74. Hardie, R.C., et al., Molecular basis of amplification in Drosophila phototransduction: roles for G protein, phospholipase C, and diacylglycerol kinase. Neuron, 2002. 36(4): p. 689–701.

## References

1. Chaudhari, A., et al., Circadian clocks, diets and aging. Nutr Healthy Aging, 2017. 4(2): p. 101–112.

2. Sato, S., et al., Circadian Reprogramming in the Liver Identifies Metabolic Pathways of Aging. Cell, 2017. 170(4): p. 664–677 e11.

3. Eckel-Mahan, K.L., et al., Reprogramming of the circadian clock by nutritional challenge. Cell, 2013. 155(7): p. 1464–78.

4. Cho, E., et al., AMP-Activated Protein Kinase Regulates Circadian Rhythm by Affecting CLOCK in Drosophila. J Neurosci, 2019. 39(18): p. 3537–3550.

5. Ramanathan, C., et al., mTOR signaling regulates central and peripheral circadian clock function. PLoS Genet, 2018. 14(5): p. e1007369.

6. Bae, S.A., et al., At the Interface of Lifestyle, Behavior, and Circadian Rhythms: Metabolic Implications. Front Nutr, 2019. 6: p. 132.

7. Kuintzle, R.C., et al., Circadian deep sequencing reveals stress-response genes that adopt robust rhythmic expression during aging. Nat Commun, 2017. 8: p. 14529.

8. Zhang, M., et al., Dysregulated metabolic pathways in age-related macular degeneration. Sci Rep, 2020. 10(1): p. 2464.

9. Vallee, A., et al., Circadian Rhythms in Exudative Age-Related Macular Degeneration: The Key Role of the Canonical WNT/beta-Catenin Pathway. Int J Mol Sci, 2020. 21(3).

10. Baba, K. and G. Tosini, Aging Alters Circadian Rhythms in the Mouse Eye. J Biol Rhythms, 2018. 33(4): p. 441–445.

11. Felder-Schmittbuhl, M.P., et al., Ocular Clocks: Adapting Mechanisms for Eye Functions and Health. Invest Ophthalmol Vis Sci, 2018. 59(12): p. 4856–4870.

12. Kawashima, M., et al., Calorie restriction (CR) and CR mimetics for the prevention and treatment of age-related eye disorders. Exp Gerontol, 2013. 48(10): p. 1096–100.

13. Baba, K., et al., The Retinal Circadian Clock and Photoreceptor Viability. Adv Exp Med Biol, 2018. 1074: p. 345–350.

14. Partch, C.L., C.B. Green, and J.S. Takahashi, Molecular architecture of the mammalian circadian clock. Trends Cell Biol, 2014. 24(2): p. 90–9.

15. Baba, K., et al., Removal of clock gene Bmal1 from the retina affects retinal development and accelerates cone photoreceptor degeneration during aging. Proc Natl Acad Sci U S A, 2018. 115(51): p. 13099–13104.

16. Sawant, O.B., et al., The Circadian Clock Gene Bmal1 Controls Thyroid Hormone-Mediated Spectral Identity and Cone Photoreceptor Function. Cell Rep, 2017. 21(3): p. 692–706.

17. Fu, Y. and K.W. Yau, Phototransduction in mouse rods and cones. Pflugers Arch, 2007. 454(5): p. 805–19.

18. Montell, C., Drosophila visual transduction. Trends Neurosci, 2012. 35(6): p. 356–63.

19. Do, M.T. and K.W. Yau, Intrinsically photosensitive retinal ganglion cells. Physiol Rev, 2010. 90(4): p. 1547–81.

20. Owens, L., et al., Effect of circadian clock gene mutations on nonvisual photoreception in the mouse. Invest Ophthalmol Vis Sci, 2012. 53(1): p. 454–60.

21. Shieh, B.H., Molecular genetics of retinal degeneration: A Drosophila perspective. Fly (Austin), 2011. 5(4): p. 356–68.

22. Nash, T.R., et al., Daily blue-light exposure shortens lifespan and causes brain neurodegeneration in Drosophila. NPJ Aging Mech Dis, 2019. 5: p. 8.

23. Baik, L.S., et al., Circadian modulation of light-evoked avoidance/attraction behavior in Drosophila. PLoS One, 2018. 13(8): p. e0201927.

24. Pittendrigh, C.S., Temporal organization: reflections of a Darwinian clock-watcher. Annu Rev Physiol, 1993. 55: p. 16–54.

25. Nippe, O.M., et al., Circadian Rhythms in Visual Responsiveness in the Behaviorally Arrhythmic Drosophila Clock Mutant Clk(Jrk). J Biol Rhythms, 2017. 32(6): p. 583–592.

26. Storch, K.F., et al., Intrinsic circadian clock of the mammalian retina: importance for retinal processing of visual information. Cell, 2007. 130(4): p. 730–741.

27. Organisciak, D.T., et al., Circadian-dependent retinal light damage in rats. Invest Ophthalmol Vis Sci, 2000. 41(12): p. 3694–701.

28. Ferrucci, L. and E. Fabbri, Inflammageing: chronic inflammation in ageing, cardiovascular disease, and frailty. Nat Rev Cardiol, 2018. 15(9): p. 505–522.

29. Fougere, B., et al., Chronic Inflammation: Accelerator of Biological Aging. J Gerontol A Biol Sci Med Sci, 2017. 72(9): p. 1218–1225.

30. Kounatidis, I., et al., NF-kappaB Immunity in the Brain Determines Fly Lifespan in Healthy Aging and Age-Related Neurodegeneration. Cell Rep, 2017. 19(4): p. 836–848.

31. Du, Y., et al., Photoreceptor cells are major contributors to diabetes-induced oxidative stress and local inflammation in the retina. Proc Natl Acad Sci U S A, 2013. 110(41): p. 16586–91.

32. Yang, Y., et al., Neuronal necrosis and spreading death in a Drosophila genetic model. Cell Death Dis, 2013. 4: p. e723.

33. Srinivasan, N., et al., Actin is an evolutionarily-conserved damage-associated molecular pattern that signals tissue injury in Drosophila melanogaster. Elife, 2016. 5.

34. Kondratov, R.V., et al., Early aging and age-related pathologies in mice deficient in BMAL1, the core componentof the circadian clock. Genes Dev, 2006. 20(14): p. 1868–73.

35. Boomgarden, A.C., et al., Chronic circadian misalignment results in reduced longevity and large-scale changes in gene expression in Drosophila. BMC Genomics, 2019. 20(1): p. 14.

36. Mazzotti, D.R., et al., Human longevity is associated with regular sleep patterns, maintenance of slow wave sleep, and favorable lipid profile. Front Aging Neurosci, 2014. 6: p. 134.

37. Kumar, S., A. Mohan, and V.K. Sharma, Circadian dysfunction reduces lifespan in Drosophila melanogaster. Chronobiol Int, 2005. 22(4): p. 641–53.

38. Libert, S., et al., Deviation of innate circadian period from 24 h reduces longevity in mice. Aging Cell, 2012. 11(5): p. 794–800.

39. Inokawa, H., et al., Chronic circadian misalignment accelerates immune senescence and abbreviates lifespan in mice. Sci Rep, 2020. 10(1): p. 2569.

40. Patel, S.A., et al., Circadian clocks govern calorie restriction-mediated life span extension through BMAL1-and IGF-1-dependent mechanisms. FASEB J, 2016. 30(4): p. 1634–42.

41. Shen, J. and J. Tower, Effects of light on aging and longevity. Ageing Res Rev, 2019. 53: p. 100913.

42. Hori, M., et al., Lethal effects of short-wavelength visible light on insects. Sci Rep, 2014. 4: p. 7383.

43. Katewa, S.D., et al., Peripheral Circadian Clocks Mediate Dietary Restriction-Dependent Changes in Lifespan and Fat Metabolism in Drosophila. Cell Metab, 2016. 23(1): p. 143–54.

44. McLay, L.K., M.P. Green, and T.M. Jones, Chronic exposure to dim artificial light at night decreases fecundity and adult survival in Drosophila melanogaster. J Insect Physiol, 2017. 100: p. 15–20.

45. Hughes, M.E., J.B. Hogenesch, and K. Kornacker, JTK_CYCLE: an efficient nonparametric algorithm for detecting rhythmic components in genome-scale data sets. J Biol Rhythms, 2010. 25(5): p. 372–80.

46. Love, M.I., W. Huber, and S. Anders, Moderated estimation of fold change and dispersion for RNA-seq data with DESeq2. Genome Biol, 2014. 15(12): p. 550.

47. Wijnen, H., et al., Control of daily transcript oscillations in Drosophila by light and the circadian clock. PLoS Genet, 2006. 2(3): p. e39.

48. Charlton-Perkins, M.A., et al., Multifunctional glial support by Semper cells in the Drosophila retina. PLoS Genet, 2017. 13(5): p. e1006782.

49. Abruzzi, K.C., et al., Drosophila CLOCK target gene characterization: implications for circadian tissue-specific gene expression. Genes Dev, 2011. 25(22): p. 2374–86.

